# TCF7L1 and TCF7 differentially regulate specific mouse ES cell genes in response to GSK-3 inhibition

**DOI:** 10.1101/473801

**Authors:** Steven Moreira, Caleb Seo, Enio Polena, Sujeivan Mahendram, Eloi Mercier, Alexandre Blais, Bradley W. Doble

## Abstract

The genome-wide chromatin occupancy of the TCF/LEF factors and its modulation by Wnt pathway activation remain poorly defined. Here, we describe mouse ES cell (mESC) lines expressing a single copy knock-in of the 3xFLAG epitope at the N-terminus of TCF7L1 and TCF7, the two most-highly expressed TCF/LEF factors in mESCs. TCF7L1 protein levels, detected by immunoblotting with a FLAG antibody, were much higher than TCF7 in mESCs maintained in standard serum- and LIF-supplemented medium, even in the presence of the GSK-3 inhibitor, CHIR99021 (CHIR). We used FLAG antibody-mediated ChIP-seq to determine TCF7 and TCF7L1 chromatin occupancy in mESCs cultured in standard medium with or without CHIR for 14 hours. TCF7 and TCF7L1 displayed very few overlapping ChIP peaks across the genome, with TCF7L1 binding significantly more genes than TCF7 in both culture conditions. Despite a reduction in total TCF7L1 protein after CHIR treatment, the TCF7L1 ChIP peak profiles were not uniformly attenuated. Our data demonstrate that TCF7L1 chromatin occupancy upon short-term CHIR treatment is modulated in a target-specific manner. Our findings also suggest that Wnt target genes in mESCs are not regulated by TCF/LEF switching, and TCF7L1, although often called a constitutive repressor, may serve as a transcriptional activator of certain target genes in CHIR-treated mESCs.

**Highlights:**

- ChIP and cytometry data suggest that TCF7L1 does not directly regulate mESC *Nanog* expression.
- TCF7L1 remains associated with β-catenin in the presence of CHIR99021.
- TCF7 and TCF7L1 display different chromatin occupancies in mESCs.
- TCF7L1 binding at specific genomic sites is variably altered by CHIR99021.

## Introduction

Wnt/β-catenin signaling plays important roles in the regulation of self-renewal and differentiation of embryonic stem cells (ESCs) and many somatic stem cells. However, nuclear signaling mechanisms remain unclear (1, 2). Wnt ligands stimulate the accumulation of cytosolic β-catenin, followed by its subsequent nuclear translocation. Nuclear βcatenin associates with T-cell factor/lymphoid enhancer factor (TCF/LEF) transcription factors bound to Wnt-responsive elements (WRE), thereby activating transcription of Wnt target genes (3). In the absence of Wnt pathway activation, glycogen synthase kinase-3 (GSK-3), as part of a multi-protein destruction complex, phosphorylates β-catenin, priming it for proteasomal degradation (4). GSK-3 inhibition promotes self-renewal and pluripotency in mouse ESCs (mESCs) (5). Ablation of GSK-3α and GSK-3β in mESCs leads to elevated levels of β-catenin and pluripotency-associated factors, OCT4 and NANOG, and highly attenuated differentiation (6, 7). mESCs can also be propagated in the presence of two inhibitors, CHIR99021 (CHIR), a GSK-3 inhibitor and activator of the Wnt/β-catenin pathway, and PD0325901, a MEK (MAP kinase kinase) inhibitor, which together, promote the homogeneous expression of a circuit of pluripotency-associated transcription factors (8). TCF7L1 and TCF7 are the most abundantly expressed TCF/LEF factors in mESCs (9), although all four family members: TCF7, TCF7L2, TCF7L1, and LEF1, are detectable at the protein level (7, 10). In mESCs, TCF7L1 was first implicated as a negative regulator of pluripotency, where it was linked to repression of *Nanog* expression (9). TCF7L1 has been thought to function as a constitutive repressor in the circuitry of transcription factors regulating self-renewal and pluripotency (11–13), although it has been suggested that Wnt stimulation may promote activation of pluripotency-associated genes by β-catenin binding to genes occupied by TCF7L1 (11).

Derepression of TCF7L1 has been demonstrated to be essential for the self-renewal enhancing effects of the GSK-3 inhibitor CHIR (14, 15), and *ESRRB* has been shown to be a critical downstream target of derepressed TCF7L1 (16). Mechanistically, it has been suggested that this derepression involves removal of TCF7L1 from the chromatin, which is mediated by its binding to β-catenin and subsequent degradation (17). TCF7L1 downregulation in response to CHIR also occurs at the transcriptional level, facilitated by c-Myc repression (18, 19). Liberation of TCF7L1 from WREs has been suggested to allow for a TCF switch, in which repressive TCF7L1 is exchanged with activating TCF7-β-catenin complexes (15). However, such a TCF switch is not required for maintenance of self-renewing mESCs or their transition from primed to naïve pluripotency (18, 20). It has been difficult to quantitatively compare chromatin occupancy of the various TCF/LEFs due to inherent differences in TCF/LEF antibody affinities and specificities. A quantitative comparison of the genomic distribution between ectopically expressed HA-tagged human TCF7 and TCF7L2 at select Wnt target genes: Axin2, Cdx1, Cdx2, T, and Sp5, demonstrated differential binding of the two factors (10). Global genomic occupancy of TCF7 and TCF7L1 in mESCs has been obtained by using chromatin immunoprecipitation followed by next-generation sequencing (ChIP-seq), which revealed that both factors regulate context-dependent Wnt signaling responses by binding to distinct target genes (21). To increase our understanding of individualized TCF/LEF functions, we developed mESC lines in which TCF7 and TCF7L1 were each 3xFLAG-tagged, allowing for quantitative comparisons. We performed ChIP-seq to identify genomic regions occupied by tagged TCF7 and/or TCF7L1 in mESCs cultured in standard medium containing serum and LIF for fourteen hours in the presence or absence of CHIR. Our data reveal very few regions of overlap between the genomic loci bound by TCF7 and TCF7L1, with TCF7L1 bound at significantly more genomic locations than TCF7 in mESCs with active or inhibited GSK-3 (low vs. high levels of nuclear β-catenin, respectively).

## Results

### A. Generation of mESC lines with N-terminal 3xFLAG tags knocked into single copies of endogenous *TCF7* and *TCF7L1* loci

To circumvent the differing affinities and specificities of commercially available TCF7 and TCF7L1 antibodies, we introduced the 3xFLAG tag into one copy of the endogenous loci of either *TCF7* or *TCF7L1* in independent wildtype (WT) mESC cell lines, by using TALEN-facilitated homologous recombination. The expression of 3xFLAG-TCF7 and 3xFLAG-TCF7L1 in targeted mESC lines was confirmed and compared by western blot analysis after 48 hours of growth in standard medium containing serum and LIF as well as medium lacking LIF (Figures 1A and S1A).

**Fig. 1.**
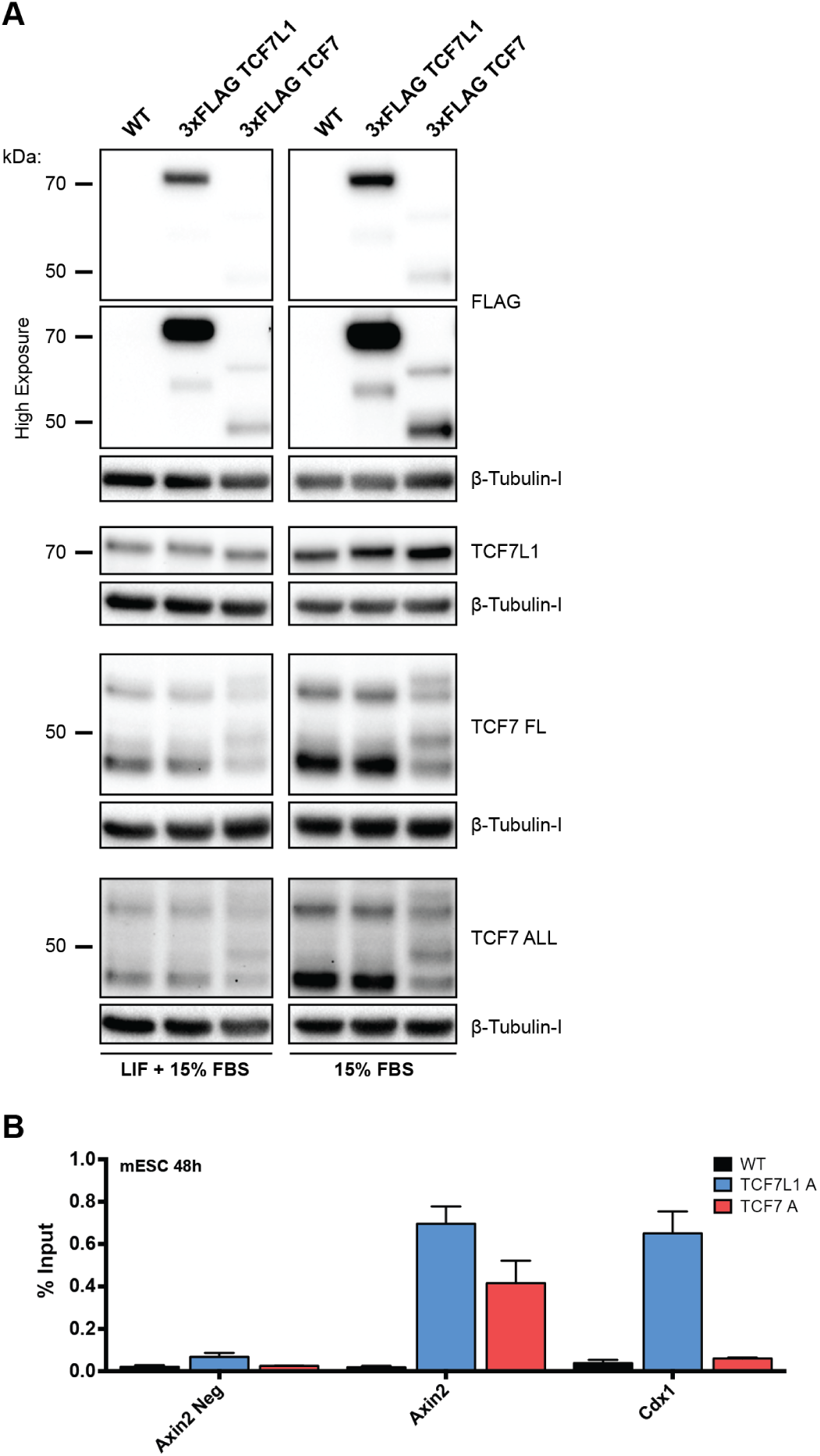
Characterization of 3xFLAG-TCF7 and 3xFLAG-TCF7L1 mESC lines. (A) Western blot analyses of 3xFLAG-TCF7 and 3xFLAG-TCF7L1 protein expression in cells maintained 48h in medium with or without LIF supplementation. Lysates were probed with antibodies against TCF7L1, TCF7, FLAG and β-Tubulin, as indicated. (B) Quantitative ChIP results obtained by using a FLAG antibody with chromatin isolated from WT, 3xFLAG-TCF7L1 and 3xFLAG-TCF7 mESCs, cultured in standard medium containing 15%FBS and LIF, for 48 h. Percent input was calculated for regions bound by TCF/LEFs in *Axin2* and *Cdx1* genes, as well as a negative control locus 11kb upstream of *Axin2*. Bars represent the mean of three independent experiments ±SEM.

Total protein levels of TCF7 and TCF7L1 in the knock-in cell lines were comparable to those observed in WT mESCs (Figure 1A and S1A). The protein levels of both TCF7 and TCF7L1 were increased in LIF-free conditions that promote differentiation (Figure 1A and S1A). Doublet bands, absent in WT blots, were clearly observed in western blots probed with anti-TCF7 antibodies, helping to confirm that 3xFLAG-TCF7 knock-in lines were heterozygous (Figure 1A and S1A).

Due to the higher molecular weight of TCF7L1, 3xFLAG-TCF7L1 and WT TCF7L1 co-migrated as a tight doublet. Heterozygosity was verified by using Sanger sequencing. 3xFLAG-TCF7L1 was expressed at much higher levels than 3xFLAG-TCF7 in standard (16-fold difference) and LIF-free (5-fold difference) medium (Figures 1A, S1A, and Table S1). These data are different from what is observed in TCF/LEF quadruple-knockout mESCs that have been rescued by repairing a single copy of endogenous *TCF7* or *TCF7L1* by using a homologous recombination strategy that introduces a 3xFLAG at the N-terminus of each protein, highlighting the importance of cross-regulation among the TCF/LEF factors (18).

To evaluate occupancy of the 3xFLAG-tagged TCF7 and TCF7L1 proteins at known Wnt target genes, *Axin2* and *Cdx1*, we used chromatin immunoprecipitation (ChIP) with a well-validated highly specific FLAG antibody, followed by quantitative RT-PCR with chromatin from mESCs cultured in standard medium for 48 hours (Figure 1B, S1B). 3xFLAG-TCF7L1 binding was detected near the transcriptional start sites of *Axin2* and *Cdx1*, as previously reported (Fig. 1B, S1B) (10, 17). Levels of 3xFLAG-TCF7L1 enrichment were only 1.67 times higher than 3xFLAG-TCF7 at the transcriptional start site of *Axin2*, despite significantly lower TCF7 protein levels (Figure 1B, S1B). We did not detect TCF7 binding at the transcriptional start site of *Cdx1*, which has been reported previously (10).

### B. Neither 3xFLAG-TCF7 nor 3xFLAG-TCF7L1 protein levels inversely correlate with NANOG protein expression in mESCs

We examined the protein expression of 3xFLAG-TCF7L1, 3xFLAG-TCF7, and NANOG in self-renewing or differentiating mESCs by using intracellular flow cytometry to assess FLAG and NANOG levels in cells grown continuously in LIF-supplemented medium for 72 hours or cells grown in LIF-free medium for 24, 48 or 72 hours (Figure 2A). In standard LIF-supplemented medium, 3xFLAG-TCF7L1 was highly expressed, with approximately 35% of the population demonstrating detectable levels, whereas 3xFLAG-TCF7 was expressed in about 14% of the population (Figures 2A, S1A, 2B, and S2B). Neither 3xFLAG-TCF7L1 nor 3xFLAG-TCF7 levels negatively correlated with NANOG levels, as evidenced by representative flow cytometric data (Figure 2A, S1A). The FLAG+ NANOG+ double-positive cells were fewer than FLAG-NANOG+ single-positive cells at days 0-2 of differentiation, but by day three the numbers of both populations were similar. Importantly, the FLAG+ NANOG+ populations displayed a higher median fluorescence intensity of NANOG than the FLAG-NANOG+ populations for both the 3xFLAG-TCF7L1 and 3xFLAG-TCF7 mESCs at all differentiation time points (Figures 2B, 2C, S2C).

**Fig. 2.**
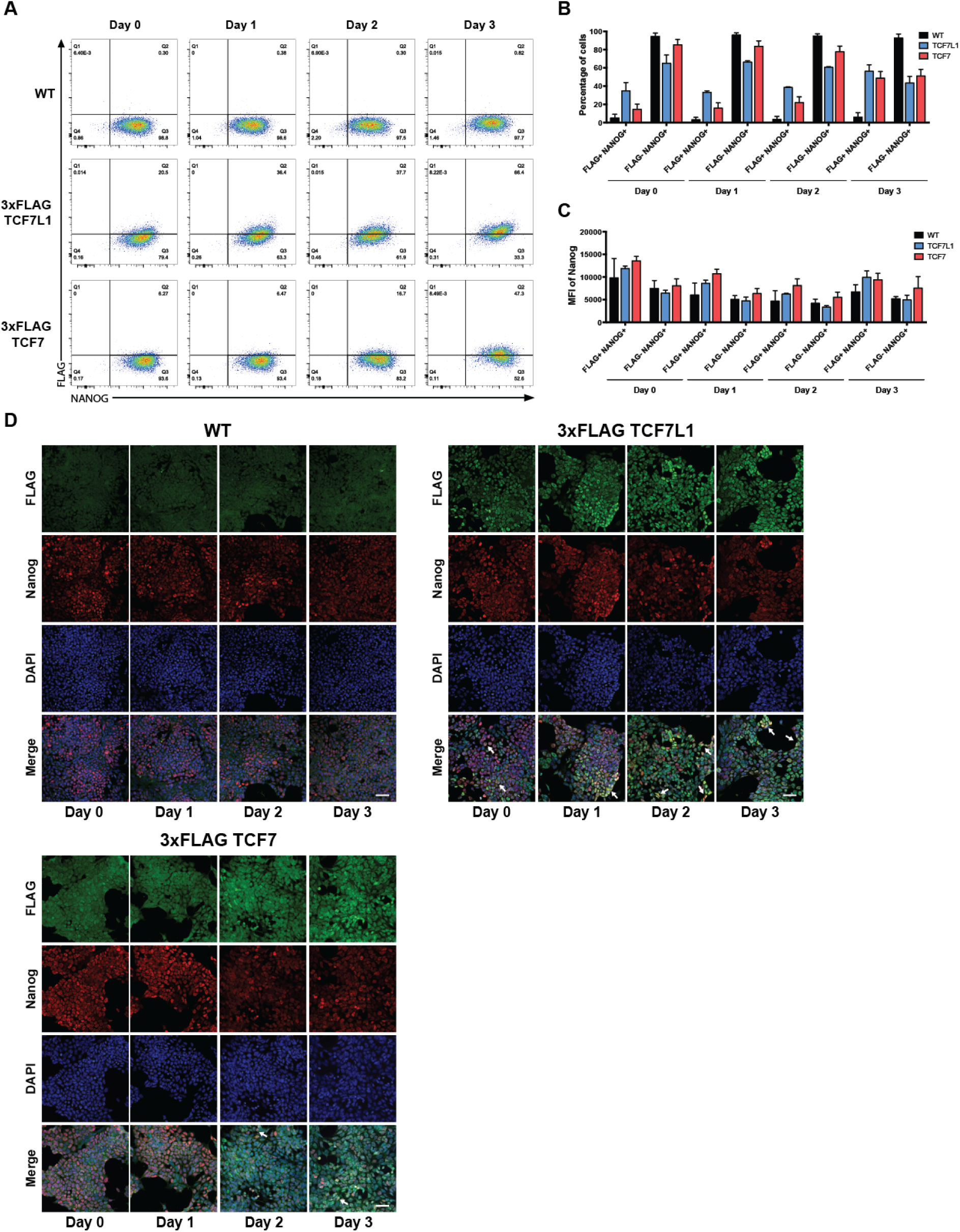
Neither TCF7L1 nor TCF7 expression inversely correlate with NANOG levels in self-renewing and differentiating mESCs. (A) Representative intracellular flow cytometric analysis of NANOG and FLAG levels in WT, 3xFLAG-TCF7L1 and 3xFLAG-TCF7 mESCs cultured in medium containing LIF (Day 0) and then differentiated in the absence of LIF (Day 1-3), as indicated. (B) Graph of the proportion of single-positive (FLAG-NANOG+) and double-positive (FLAG+ NANOG+) cells. Bars represent the mean of three independent experiments ±SEM. (C) Graph of the median fluorescence intensity of NANOG in single-positive (FLAG-NANOG+) and double-positive (FLAG+ NANOG+) cells. Bars represent the mean of three independent experiments ±SEM. (D) Immunofluorescence analysis of WT, 3xFLAG-TCF7L1 and 3xFLAG-TCF7 mESCs, cultured in medium containing LIF (Day 0) and then differentiated in the absence of LIF (Day 1-3), as indicated. Cells were stained for NANOG, FLAG and DAPI. Scale bar represents 50 µm. White arrows indicate cells with high levels of FLAG and NANOG.

Immunofluorescence microscopy of mESCs treated in the same manner as those analyzed by flow cytometry supported our flow cytometric analyses (Figure 2D, S1D). TCF7L1 expression, assessed with a FLAG antibody, revealed a heterogeneous nuclear expression pattern, with fluorescence increasing throughout the duration of LIF removal and peaking at 72 hours (Fig. 2D, S1D). Similarly, we observed heterogeneous nuclear and diffuse cytosolic expression of TCF7, which also increased throughout the course of LIF withdrawal, peaking again at 72 hours (Figure 2D, S1D). We also observed heterogeneous nuclear NANOG staining, which was highest at Day 0, before LIF was withdrawn from the culture medium (Figure 2D, S1D). Unexpectedly, many mESCs with elevated levels of NANOG and TCF7L1 persisted throughout the differentiation time course (Fig. 2D, S1D). We also observed a small number of mESCs with elevated levels of NANOG and TCF7 at 48 and 72 hours after LIF withdrawal (Figure 2D, S1D).

### C. Consequences of GSK-3 inhibition on TCF7L1 and TCF7 function in mESCs

Derepression of TCF7L1 has been described as being essential for reinforcing the pluripotent state in mESCs treated with CHIR (14, 15, 22). We sought to explore the mechanisms underlying this derepression with our unique system. To determine the timepoint at which maximal Wnt pathway activation occurred in response to CHIR, we used a lentiviral TCF/LEF reporter system in wildtype mESCs. The fluorescent protein, mCherry, driven by a constitutive promoter identifies transduced cells, whereas GFP expression is driven by tandem Wnt-responsive elements and is only expressed in the presence of transactivating β-catenin–TCF/LEF complexes.

We used flow cytometry to determine mCherry and GFP levels in mESCs treated with 5 µM CHIR, examining treated cells every 2 hours, for 18 hours. We observed minimal GFP expression in transduced control mESCs treated with DMSO (1.12%), with nearly the entire population expressing the transduction marker, mCherry (91.5%) (Figure 3A). Over the course of CHIR treatment, there was an approximately 6% increase in GFP-positive cells at every 2-hour timepoint, until 16 hours, where levels plateaued, with the GFP-positive population stabilizing at approximately 40% (Figure 3A). Based on these results, in subsequent experiments mESCs were treated with 5 µM CHIR for 14 hours. The effects of CHIR have been shown previously to dramatically reduce TCF7L1 levels while stimulating the expression of TCF7 (17–19). We used Western blot analysis to evaluate 3xFLAG-TCF7L1 and 3xFLAG-TCF7 protein levels in cells treated with 5 µM CHIR or DMSO (vehicle), for 14-hours (Figure 3B, S3A). Consistent with previous studies, protein levels of TCF7L1 were reduced (approx. 1.4-fold), whereas levels of TCF7 increased (approx. 2.5-fold) (Fig. 3B, S3A, Table S2). TCF7L1 was expressed at higher levels than TCF7, in both DMSO (approx. 15-fold) and 5 µM CHIR (approx. 3-fold) conditions (Figure 3B, S3A, Table S2).

**Fig. 3.**
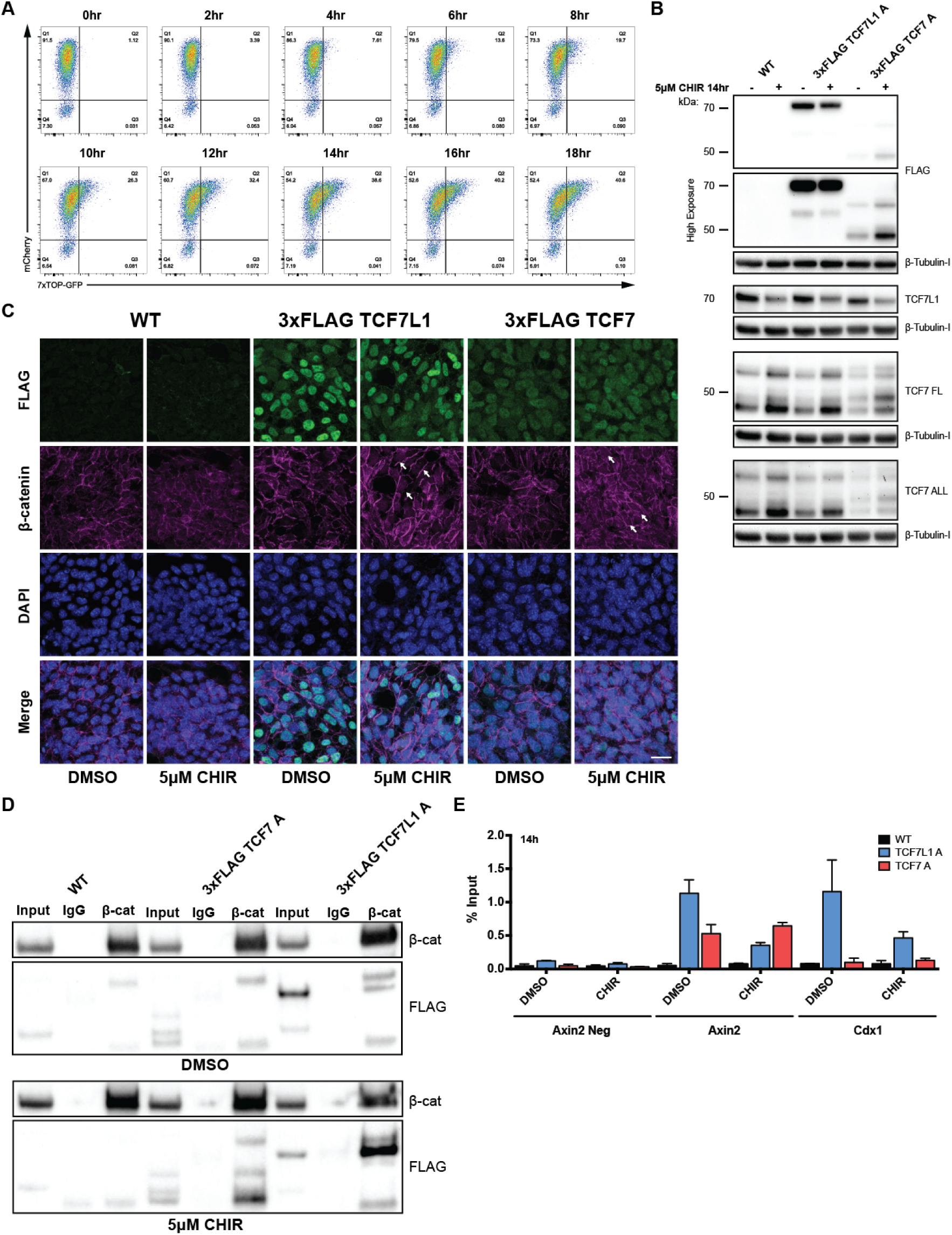
GSK-3 inhibition promotes association of β-catenin with 3xFLAG-TCF7 and 3xFLAG-TCF7L1. (A) Flow cytometric analysis of WT mESCs transduced with a GFP-based TCF-reporter cultured in standard LIF-supplemented medium and treated with 5 µM CHIR every 2 h for 18 h, as indicated. (B) Western blot analysis of WT, 3xFLAG-TCF7L1 and 3xFLAG-TCF7 mESCs, cultured in standard LIF-supplemented medium and treated with 5 µM CHIR or DMSO control for 14 h. Lysates were probed with antibodies against TCF7L1, TCF7, FLAG and β-Tubulin, as indicated. (C) Immunofluorescence analysis of WT, 3xFLAG-TCF7L1 and 3xFLAG-TCF7 mESCs, cultured in standard LIF-supplemented medium and treated with 5 µM CHIR or DMSO control for 14 h. Cells were stained for β-catenin, FLAG and DAPI. Scale bar represents 20 µm. White arrows indicate cells with elevated levels of FLAG and nuclear localization of β-catenin. (D) Co-immunoprecipitation analysis of WT, 3xFLAG-TCF7L1 and 3xFLAG-TCF7 mESCs, cultured in standard LIF-supplemented medium and treated with 5 µM CHIR or DMSO control for 14h. Lysates were immunoprecipitated using β-catenin antibody and were probed with antibodies against β-catenin and FLAG, as indicated. (E) Quantitative ChIP using the FLAG antibody on WT, 3xFLAG-TCF7L1 and 3xFLAG-TCF7 mESCs, cultured in standard LIF-supplemented medium and treated with 5 µM CHIR or DMSO control for 14 h. Percent input was calculated for WREs in *Axin2* and *Cdx1*, as well as a negative control locus 11kb upstream of *Axin2*. Bars represent the mean of three independent experiments ±SEM.

To examine the effects of CHIR treatment on a population of 3xFLAG-TCF7L1 and 3xFLAG-TCF7 mESCs, we also used immunofluorescence analyses (Figure 3C, S3B). Consistent with immunofluorescence performed in standard LIF-containing medium, mESCs stained for FLAG revealed heterogeneous expression of TCF7L1 and TCF7 in DMSO and CHIR conditions (Figure 3C, S3B). Although there was an overall reduction in total TCF7L1 protein levels in response to CHIR treatment (Figure 3B, S3A), we observed several cells that sustained elevated levels TCF7L1, as assessed by FLAG staining (Figure 3C, S3B). Similarly, although total TCF7 protein levels increased after CHIR treatment (Figure 3B, S3A), we observed some cells with slightly elevated levels of TCF7 in the presence of CHIR (Figure 3C, S3B). To determine whether the levels of β-catenin correlated with elevated or reduced levels of TCF7L1 and TCF7, respectively, we also performed immunofluorescence analysis of ’active’ β-catenin (Fig. 3C, S3B). We observed similar levels of nuclear active β-catenin in cells containing high and low levels of 3xFLAG-TCF7L1 or 3xFLAG-TCF7 (Figure 3C, S3B).

Intrigued by the apparent lack of a correlation between activated β-catenin and TCF7L1/TCF7 levels, we sought to determine which of the two factors associated with higher levels of β-catenin in the presence of CHIR. We analyzed the 3xFLAG-TCF/β-catenin interactions, by immunoprecipitating β-catenin and subsequently probing for 3xFLAG-TCF7L1 and 3xFLAG-TCF7 using anti-FLAG antibody, in lysates from mESCs treated with 5µM CHIR (Figure 3D). β-catenin was associated with higher levels of TCF7L1 compared to TCF7, in the presence of CHIR (Figure 3D), in agreement with our data showing higher TCF7L1 proteins levels compared to TCF7, even in the presence of CHIR (Figure 3B). A reduction in TCF7L1 occupancy at *Axin2* and *Cdx1* has previously been observed in mESCs treated with CHIR (17). We were therefore interested in comparing the effects of CHIR on the occupancy of both TCF7L1 and TCF7 at these two Wnt-associated genes. We performed ChIP using the FLAG antibody, followed by quantitative RT-PCR on 3xFLAG-TCF7L1 and 3xFLAG-TCF7 mESCs cultured in standard medium containing LIF treated with 5µM CHIR or DMSO for 14 hours (Figure 3E, S3C). Consistent with previous data, in the presence of CHIR we observed a reduction in TCF7L1 occupancy at a locus near the *Axin2* promoter, but only a slight reduction in occupancy at the Cdx1 promoter (Figure 3E, S3C). Unexpectedly, in response to CHIR, despite an increase in TCF7 protein (Figure 3B, S3A), we did not observe a substantial increase in TCF7 occupancy at the promoter of *Axin2*, nor *Cdx1* (Figure 3E, S3C). However, there was more TCF7 compared to TCF7L1, at the promoter of *Axin2* in the presence of CHIR (Figure 3E, S3C).

### D. Examining genome-wide TCF7L1 and TCF7 chromatin occupancy

Having observed differential binding of 3xFLAG-TCF7L1 and 3xFLAG-TCF7 to *Axin2* and *Cdx1* loci, we were interested in elucidating the global differences in chromatin distribution between these two TCF factors. We performed ChIP-seq analyses using the FLAG antibody, on wildtype, 3xFLAG-TCF7L1 and 3xFLAG-TCF7 mESCs cultured under standard LIF-supplemented conditions and after 14 hours of 5µM CHIR treatment. ChIP-seq reads were aligned to the mouse genome using BWA, where duplicates and low-quality reads were discarded. Subsequent peaks were identified using MACS, followed by *de novo* motif discovery and gene ontology analysis performed using HOMER. We identified 760 (in 658 genes) and 582 (in 527 genes) 3xFLAG-TCF7L1-bound regions (peaks) in standard medium and in response to CHIR treatment for 14 hours, respectively (Table S3). Among the 3xFLAG-TCF7L1-bound genes, there were 359 genes (43.46%) of overlap, in both control and CHIR conditions (Figure S4B). 299 (36.20%) and 168 (20.33%), 3xFLAG-TCF7L1-bound genes were unique to control and CHIR conditions, respectively (Figure S4B). We sought to validate the 658 genes bound by 3xFLAG-TCF7L1 with a previously published list of 1351 TCF7L1 bound genes, which was also obtained from mESCs maintained in standard media conditions with serum and LIF (Figure S4A) (12). Surprisingly, only 177 genes were shared between the two TCF7L1 ChIP-seq datasets (Figure S4A). A majority of 3xFLAG-TCF7L1 peaks, 37% in control conditions and 38% in CHIR, were bound within a gene’s exon or intron (Figure 4A TOP). 9% of control and 8% of CHIR 3xFLAG-TCF7L1 peaks were bound to ‘promoter’ regions within 2kb of a transcriptional start site (TSS) (Figure 4A TOP). Additionally, we observed 31% of 3xFLAG-TCF7L1 peaks distributed at ‘enhancer’ regions, falling within 2kb to 100kb upstream of a TSS, in both control and CHIR conditions (Figure 4A TOP). Interestingly, the genomic distribution of 3xFLAG-TCF7L1 bound regions remained nearly unaffected by CHIR treatment (Figure 4A TOP). Our 3xFLAG-TCF7 ChIP-seq identified considerably less peaks, 17 (in 16 genes) and 84 (in 75 genes), in control and CHIR conditions, respectively (Table S3). Among these 3xFLAG-TCF7 bound genes, there were 12 genes (15.19%) with overlapping peaks, in control and CHIR conditions (Figure S4B). Four (5.06%) and 63 (79.75%), 3xFLAG-TCF7 bound genes were unique to control and CHIR conditions, respectively (Figure S4B). The significantly lower number of 3xFLAG-TCF7 bound regions can likely be attributed to lower 3xFLAG-TCF7 protein levels observed in the cells, which only slightly increases upon CHIR supplementation (Figure 3B). These 3xFLAG-TCF7 bound regions are biologically relevant in this context, and they display robust 3xFLAG-TCF7 peaks, despite its low protein levels.

**Fig. 4.**
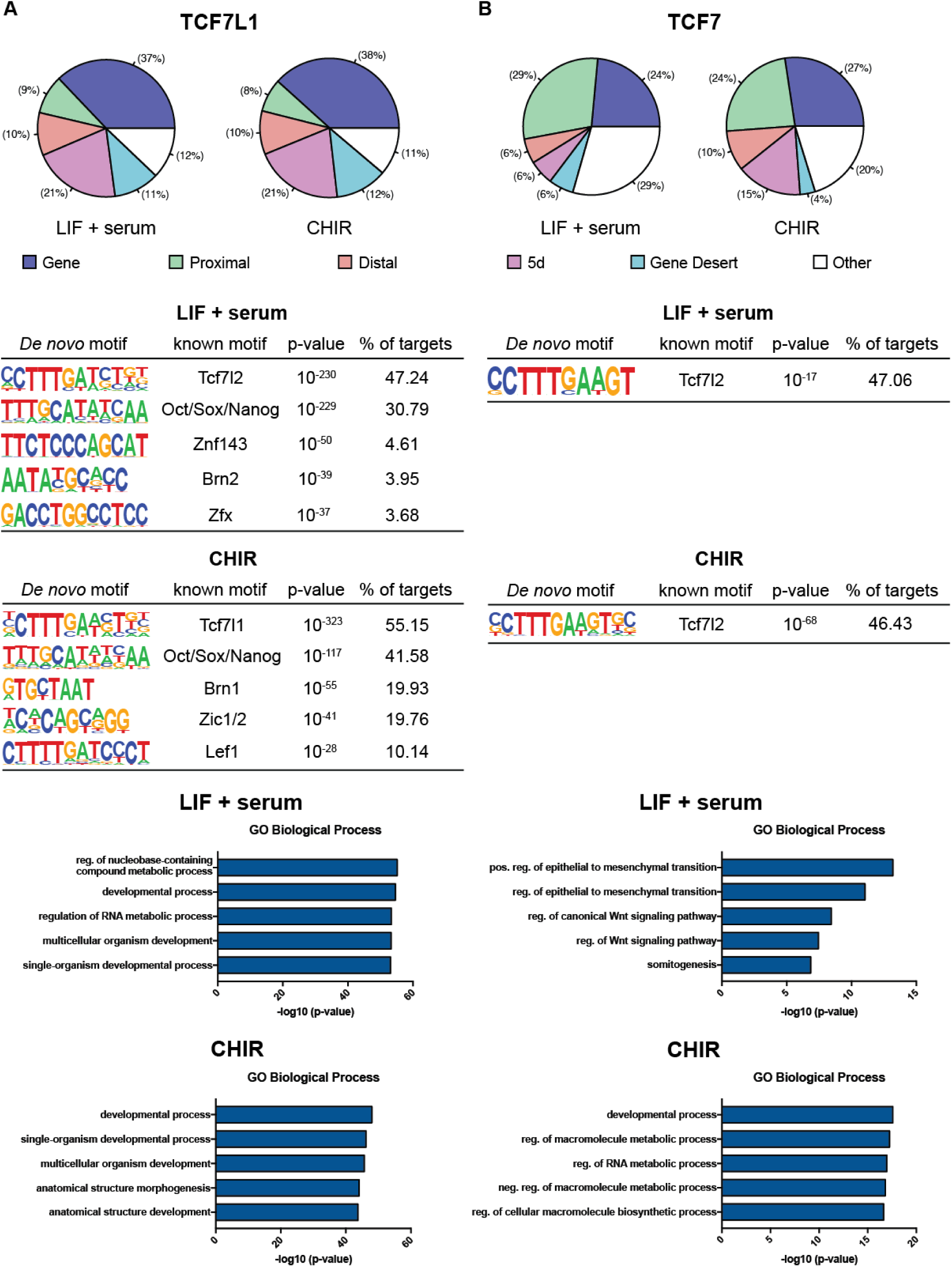
Characterization of genomic regions bound by 3xFLAG-TCF7L1 or 3xFLAG-TCF7 in mESCs. (A) TOP: Genomic distribution of 3xFLAG-TCF7L1 ChIP-seq peaks for mESCs maintained in standard medium (LIF and serum) or medium supplemented with 5µM CHIR for 14h. Gene: exon or intron. Proximal: 2kb upstream of a transcriptional start site (TSS). Distal: between 10kb upstream and 2kb upstream of a TSS. 5d: between 100kb upstream and 10kb upstream of a TSS. Gene desert: 100kb upstream or downstream of a TSS. Other: anything not included above. MID: Enriched motifs from a *de novo* motif search of sequences contained in 3xFLAG-TCF7L1 peaks in control and CHIR conditions. BOT: Gene ontology analysis of 3xFLAG-TCF7L1 ChIP-seq peaks in control and CHIR conditions. Values are presented as negative log-base 10 of their p-values. (B) TOP: Genomic distribution of 3xFLAG-TCF7 ChIP-seq peaks in control and CHIR conditions. Gene: exon or intron. Proximal: 2kb upstream of a TSS. Distal: between 10kb upstream and 2kb upstream of a TSS. 5d: between 100kb upstream and 10kb upstream of a TSS. Gene desert: 100kb upstream or downstream of a TSS. Other: anything not included above. MID: Enriched motifs from a *de novo* motif search of sequences contained in 3xFLAG-TCF7 peaks in control and CHIR conditions. BOT: Gene ontology analysis of 3xFLAG-TCF7 ChIP-seq peaks in control and CHIR conditions. Values are presented as negative log-base 10 of their p-values.

We also assessed the genomic distribution of 3xFLAG-TCF7 peaks and found that the majority, 24% in control and 27% in CHIR conditions, were bound within a gene’s exons or introns (Figure 4B TOP). A large portion were also bound to ‘promoter’ regions within 2 kb of a TSS, 29% and 24%, in control and CHIR conditions, respectively (Figure 4B TOP). Furthermore, we observed that 12% and 25% of 3xFLAG-TCF7 peaks, in control and CHIR conditions, respectively, were distributed at ‘enhancer’ regions, falling within 2 kb to 100 kb upstream of a TSS (Figure 4B TOP). Interestingly, 29% and 20% of 3xFLAG-TCF7 peaks, in control and CHIR conditions, respectively, were located at regions that could not be categorized (Figure 4B TOP).

Due to our ability to utilize the FLAG antibody to perform ChIP-seq on both 3xFLAG-tagged TCF7L1 and 3xFLAG-TCF7, we were interested in comparing the genes bound by both factors. Intriguingly, we did not observe a large overlap in the genes bound by both TCF7L1 or TCF7 (Figure S4C). In control medium, 6 genes were common between both factors, which represents 37.5% and 0.92% of genes bound by 3xFLAG-TCF7 and 3xFLAG-TCF7L1, respectively (Figure S4C). We observed a similar trend in the presence of CHIR, with 33 genes bound by both factors, representing 44% and 6.68% of genes bound by TCF7 and TCF7L1, respectively (Figure S4C).

Importantly, the TCF/LEF consensus binding motif (Wnt responsive element; WRE) was the top *de novo* motif identified, representing 47.24% and 55.15% of 3xFLAG-TCF7L1 peaks, in control and CHIR conditions, respectively (Figure 4A MID). An additional TCF/LEF consensus binding motif was discovered in CHIR, representing 10.14% of 3xFLAG-TCF7L1 peaks (Figure 4A MID). As anticipated, the top *de novo* motif observed in 3xFLAG-TCF7 peaks was also the TCF/LEF consensus binding motif, representing 47.06% and 46.43% of TCF7 peaks, in control and CHIR conditions, respectively (Figure 4B MID).

In addition to the TCF/LEF consensus binding motif, 3xFLAG-TCF7 and 3xFLAG-TCF7L2 are capable of binding to adjacent ‘helper sites’ via a cysteine-clamp (C-clamp) domain,(23, 24). To identify helper sites, we scanned 3xFLAG-TCF7 and 3xFLAG-TCF7L1 for the presence of WRE motif peaks using HOMER, followed by scanning for helper sites located 200 bp upstream or downstream of the WRE. As anticipated, a considerable fraction of 3xFLAG-TCF7 peaks contained both a WRE and the helper site, specifically, 12 (70.59%) and 40 (47.62%) peaks, in control or CHIR conditions, respectively (Table S4). By contrast, 3xFLAG-TCF7L1 peaks were not enriched for helper sites, as only 66 (11.34%) and 72 (9.47%) peaks identified in mESC or CHIR conditions, respectively, contained both a WRE and helper site, consistent with the absence of a c-clamp in TCF7L1 (Table S4).

TCF7L1 is a part of an extended network of transcription factors regulating pluripotency. This was reflected in our ChIP-seq data, which revealed a *de novo* motif associated with POU5F1/SOX2/NANOG co-bound genomic regions (Figure 4A MID) (25). This motif was the second most enriched *de novo* motif, which was present in 30.79% of 3xFLAG-TCF7L1 peaks in control conditions and surprisingly, represented 41.58% of TCF7L1 peaks in the presence of CHIR (Figure 4A MID). *De novo* motifs associated with *Znf143*, *Brn2*, and *Zfx* were also discovered in control conditions, although these motifs were present in less than 5% of 3xFLAG-TCF7L1 peaks (Figure 4A MID). Intriguingly, in CHIR conditions, *de novo* motifs associated with *Brn1* and *Zic1/2*, were discovered in 19.93% and 19.76% of 3xFLAG-TCF7L1 associated peaks (Figure 4A MID).

Gene Ontology (GO) analysis was performed using HOMER to search for functional biological process GO categories enriched in 3xFLAG-TCF7L1 bound genes (Figure 4A BOT). Three out of the top five most enriched biological process GO terms for 3xFLAG-TCF7L1 bound genes in both control and CHIR conditions were associated with developmental processes (Figure 4A BOT). In control conditions, 3xFLAG-TCF7L1-bound genes were also associated with the biological process GO terms ’regulation of nucleobase-containing compound metabolism’ and ’regulation of RNA metabolism’ (Figure 4A BOT). Finally, in CHIR conditions, the biological process GO terms ’anatomical structure morphogenesis’ and ’anatomical structure development’ were also enriched in regions bound by 3xFLAG-TCF7L1 (Figure 4A BOT).

Gene ontology analysis of genes bound by 3xFLAG-TCF7 was also conducted (Figure 4B BOT). In control conditions, the two most enriched biological process GO terms for 3xFLAG-TCF7 peaks, were both associated with regulation of the epithelial to mesenchymal transition (Figure 4B BOT). Furthermore, two biological process GO terms associated with regulation of Wnt signaling and canonical Wnt signaling were identified, despite only identifying 16 genes bound by 3xFLAG-TCF7 in control conditions (Figure 4B BOT). In CHIR-treated mESCs, three out of the top five most enriched biological process GO terms for 3xFLAG-TCF7 bound genes, were associated with macromolecule metabolism or biosynthesis (Figure 4B BOT). The remaining two biological process GO terms ’developmental process’ and ’regulation of RNA metabolism’ were also observed in genes bound by 3xFLAG-TCF7L1 in control conditions(Figure 4B BOT).

### E. 3xFLAG-TCF7L1 and 3xFLAG-TCF7 display differential chromatin occupancy

To further explore the differential binding of 3xFLAG-TCF7L1 and 3xFLAG-TCF7 to their respective target genes and the effects of Wnt pathway activation using CHIR, we employed CSAW (26). CSAW allows for quantitative evaluation of differential binding between experimental conditions and samples instead of qualitatively assessing absence or presence (26). Additionally, CSAW allows for rigorous statistical analysis, controlling for false discovery rate across constant regions or windows across the genome and accounting for biological variation between samples (26).

We used CSAW with the 3xFLAG-TCF7L1 and 3xFLAG-TCF7 ChIP-seq datasets to obtain differential binding data, comparing not only genes bound by a single factor in both conditions, but also comparing genes bound by both factors in a single condition (Table S5). We first analyzed 3xFLAG-TCF7L1 occupancy in control vs CHIR conditions (Figure 5A). Surprisingly, 307 peaks in genes including *Bmper*, *Bdh1*, *Ext1*, and *Dkk4*, were preferentially bound in CHIR, compared to 332 preferentially bound peaks identified in control conditions, found in genes such as *Myb*, *Ptch1*, *Fgfr2*, and *Trp53bp1* (Figure 5A). Differential binding analysis performed on 3xFLAG-TCF7, revealed 17 peaks preferentially bound in control conditions (Figure 5A). However, it should be noted that with the exception of *Camk2a*, none of these genes were identified as peaks with MACS (Figure 5A). Conversely, in CHIR-treated mESCs, 463 peaks were preferentially bound, including peaks found in the genes, *Tcf7l2*, *Gbx2*, *Bmper*, and *Mllt6* (Figure 5A). To determine regions differentially bound by 3xFLAG-TCF7L1 and 3xFLAG-TCF7, we also compared the binding of the two factors in a single condition. In control conditions, 40 and 2036 regions were preferentially bound by 3xFLAG-TCF7 and 3xFLAG-TCF7L1, respectively (Figure 5B). Of the 40 regions preferentially bound by 3xFLAG-TCF7, a majority of those topmost enriched were associated with genes involved in Wnt signaling, namely *Zfp703*, *Tcf7*, *Nkd1*, *Lef1*, *Lgr5*, and *Ccnd1* (Figure 5B). Similarly, the most enriched genes preferentially bound by 3xFLAG-TCF7L1 were associated with both Wnt signaling and pluripotency, and included: *Mllt6*, *Ctnnd2*, *Id3*, *Cdx1*, *Klf2*, *Sp5* and *Tfcp2l1* (Figure 5B).

**Fig. 5.**
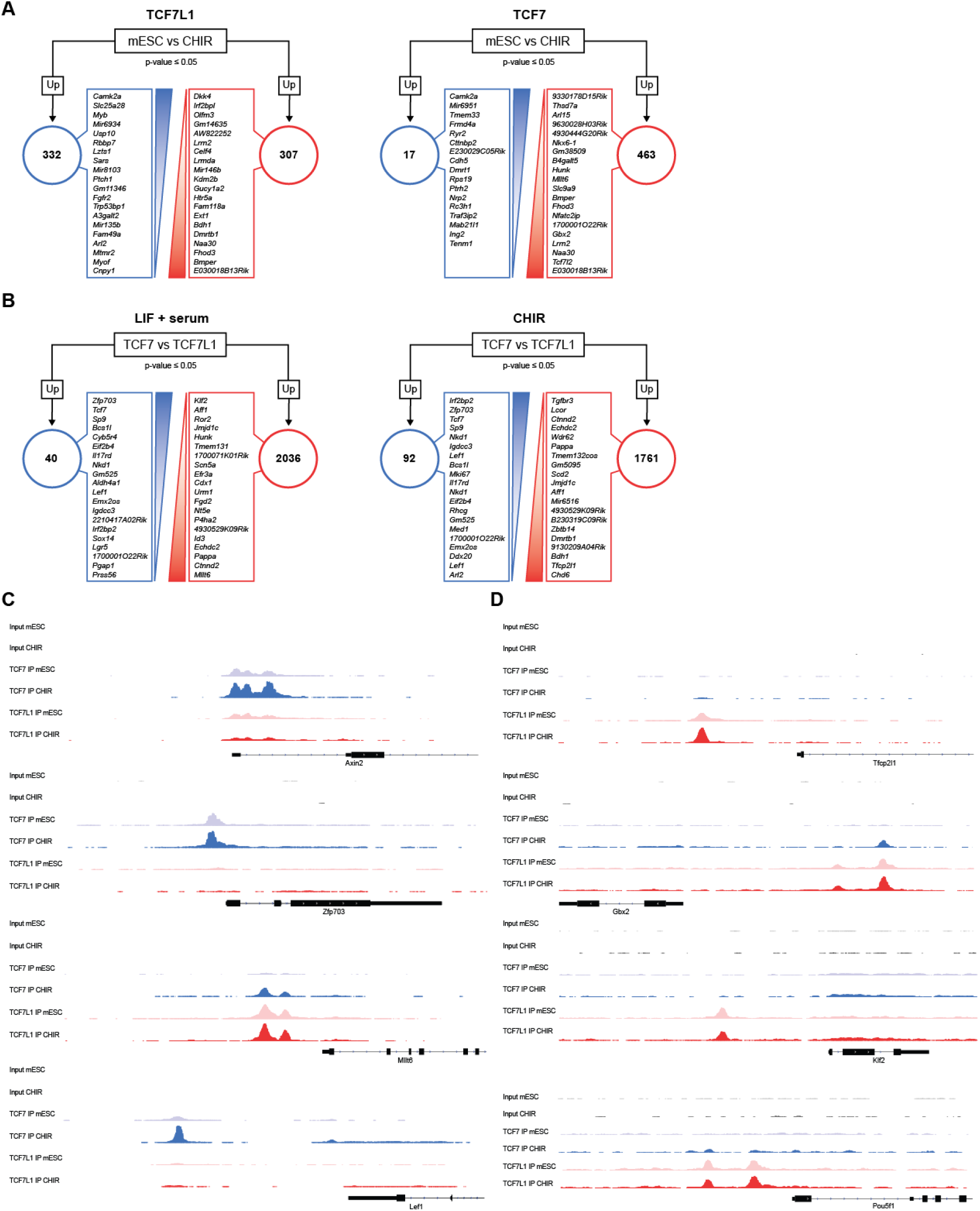
3xFLAG-TCF7L1 and 3xFLAG-TCF7 exhibit differential occupancy at Wnt target and pluripotency-associated genes. (A) Target genes preferentially bound by 3xFLAG-TCF7L1 (LEFT) or 3xFLAG-TCF7 (RIGHT) in control and CHIR conditions. Differential binding was determined using CSAW with a cutoff of 0.05. (B) Target genes preferentially bound by 3xFLAG-TCF7 versus 3xFLAG-TCF7L1, in mESCs cultured control (LEFT) or CHIR conditions (RIGHT). Differential binding was determined by using CSAW with a cutoff of 0.05. (C) Genomic tracks showing 3xFLAG-TCF7 (blue) and 3xFLAG-TCF7L1 (red) peaks at selected Wnt-associated genes in mESCs cultured in control medium (light shade) or CHIR-containing medium (dark shade). Pooled control and CHIR input peaks are in grey and dark grey, respectively. Genomic positions reflect NCBI mouse genome build mm10. (D) Genomic tracks showing 3xFLAG-TCF7 (blue) and 3xFLAG-TCF7L1 (red) peaks at selected pluripotency-associated genes in mESCs cultured control (light shade) or CHIR conditions (dark shade). Genomic tracks showing pooled control and CHIR input peaks are in grey and dark grey, respectively. Genomic positions reflect NCBI mouse genome build mm10.

Conversely, in the presence of CHIR, 92 and 1761 regions were preferentially bound by 3xFLAG-TCF7 and 3xFLAG-TCF7L1, respectively (Figure 5B). Of the 92 regions preferentially bound by 3xFLAG-TCF7, the topmost abundant genes were once again associated with genes involved in Wnt signaling, such as *Zfp703*, *Tcf7*, *T*, *Nkd1*, *Lef1*, *Axin2*, and *Ccnd1* (Figure5B). Similarly, a majority of the most enriched genes preferentially bound by 3xFLAG-TCF7L1 were associated with both Wnt signaling and pluripotency, and included: *Tfcp2l1*, *Bdh1*, *Ctnnd2*, *Sp5*, *Fzd7*, *Dkk4* and *Lifr* (Figure 5B).

To further explore 3xFLAG-TCF7L1 and 3xFLAG-TCF7 binding patterns, we examined the peak profiles at select Wnt and pluripotency-associated genes. In both conditions, 3xFLAG-TCF7 was associated with the Wnt related genes, *Zfp703*, *Lef1*, *Msx2*, *Tcf7*, *Lef1*, and *Nkd1* (Figure 5C, S5A). All of these genes were bound by 3xFLAG-TCF7 in control conditions and demonstrated increased binding after addition of CHIR for 14 hours (Figure 5C, S5A). Moreover, in the presence of CHIR, 3xFLAG-TCF7 was detected at the Wnt target, *T* (Figure S5A). By contrast, 3xFLAG-TCF7L1 was located at only two Wnt-associated genes, *Lgr5* and *Dkk4* (Figure S5A). *Lgr5* was bound by 3xFLAG-TCF7L1 in both control and CHIR conditions, however a slight reduction in binding was observed in response to CHIR (Figure 5C, S5A). Notably, the site bound by 3xFLAG-TCF7L1 in intron 8 of the *Lgr5* gene was not bound by 3xFLAG-TCF7, although 3xFLAG-TCF7 bound an alternate site in the promoter region of *Lgr5* (Figure S5A). 3xFLAG-TCF7L1 was detected in the promoter region of *Dkk4*, but only in the presence of CHIR (Figure S5A), wherease no 3xFLAG-TCF7 ChIP signal was detected in the *Dkk4* locus in either condition. In control conditions, TCF7L1 was also predominantly detected at *Mllt6*, *Sp5*, *Cdx1*, and *Tcf7l1* (Figure 5C, S5A).

Minimal overlap in 3xFLAG-TCF7L1 and 3xFLAG-TCF7 occupancies at Wnt-associated genes was observed in control conditions, with the exception of *Axin2* and *Lef1*. At *Axin2*, both factors were equally bound within the same regions, located near the 5’UTR (Figure 5C). Treating with CHIR led to a reduction in 3xFLAG-TCF7L1 binding, whereas 3xFLAG-TCF7 binding was increased, at *Axin2*, conflicting with our previous qChIP results (Figure 5C, 3E). However, mESCs utilized for ChIP-seq were cultured on feeders beforehand for two passages, whereas the mESCs used for quantitative ChIP had been maintained for multiple passages on gelatin-coated plates, which may have altered cellular properties such as chromatin accessibility. This likely also effected *Cdx1* occupancy as assessed by ChIP-seq, as we were able to detect 3xFLAG-TCF7 in the presence of CHIR (Figure S5A). Furthermore, at *Cdx1*, we observed an increase, not a decrease, in 3xFLAG-TCF7L1 binding in response to CHIR stimulation (Figure S5A). At *Lef1*, 3xFLAG-TCF7 was bound to a higher extent than 3xFLAG-TCF7L1 in both control and CHIR conditions (Figure 5C). In the presence of CHIR, 3xFLAG-TCF7L1 and 3xFLAG-TCF7 demonstrated some overlap with respect to Wnt related genes. In addition to *Axin2* and *Cdx1*, both factors were detected at *Mllt6*, *Sp5*, *Tcf7l2*, and *Tcf7l1* (Figure 5C, S5A), although 3xFLAG-TCF7L1 was more abundant at *Mllt6*, *Tcf7l2* and *Tcf7l1* in the CHIR condition (Figure 5C, S5A).

3xFLAG-TCF7L1 was predominantly associated with pluripotency-related genes, occupying regions within: *Tfcp2l1*, *Gbx2*, *Klf2*, *Pou5f1*, *Klf4*, *Sox2*, *Nanog*, *Esrrb*, *Nodal*, *Zfp42*, and *Dppa3*, in both control and CHIR conditions (Figure 5D, S5B). Unexpectedly, 3xFLAG-TCF7L1 occupancy was only slightly reduced in response to CHIR at *Klf2*, *Nanog* and one of the three peaks discovered at both *Klf4* or *Esrrb* (Figure 5D, S5B). Furthermore, 3xFLAG-TCF7L1 occupancy at *Tfcp2l1*, *Gbx2*, *Pou5f1*, *Sox2*, *Nodal*, *Zfp42*, and two of the three peaks identified at both *Klf4* or *Esrrb*, remained unchanged or actually increased upon CHIR treatment (Figure 5D, S5B). 3xFLAG-TCF7, in control medium, was only slightly enriched at *Sox2* and treatment with CHIR led to an increase in 3xFLAG-TCF7 binding (Figure S5B). At *Sox2*, both 3xFLAG-TCF7L1 and 3xFLAG-TCF7 were identified at similar levels (Figure S5B). In the presence of CHIR, we observed slight increases in 3xFLAG-TCF7 binding to *Gbx2*, *Pou5f1*, and *Nodal*, although 3xFLAG-TCF7L1 levels remained higher at all three gene loci (Figure 5D, S5B).

### F. TCF7L1 acts as a transcriptional activator in response to CHIR-mediated GSK-3 inhibition

*Id3* was one of the most significantly upregulated genes in cells lacking all full-length TCF/LEF factors (18). As we observed that 3xFLAG-TCF7L1 remained bound to pluripotency-associated genes in the presence of CHIR, we analyzed the 3xFLAG-TCF7L1 binding profile at the *Id3* locus. In control conditions, we observed a significant 3xFLAG-TCF7L1 peak located 1.5 kb downstream of the *Id3* gene (Figure 6A). CHIR treatment for 14 hours did not cause apparent 3xFLAG-TCF7L1 dissociation from the chromatin. Instead, the 3xFLAG-TCF7L1 peak at *Id3* was detected to an equal or slightly greater extent (Figure 6A). Additionally, only a negligible amount of 3xFLAG-TCF7 was bound to the *Id3*associated peak in the presence of CHIR (Figure 6A). This suggested to us that CHIR could potentially convert TCF7L1 from a transcriptional repressor into a transcriptional activator.

**Fig. 6.**
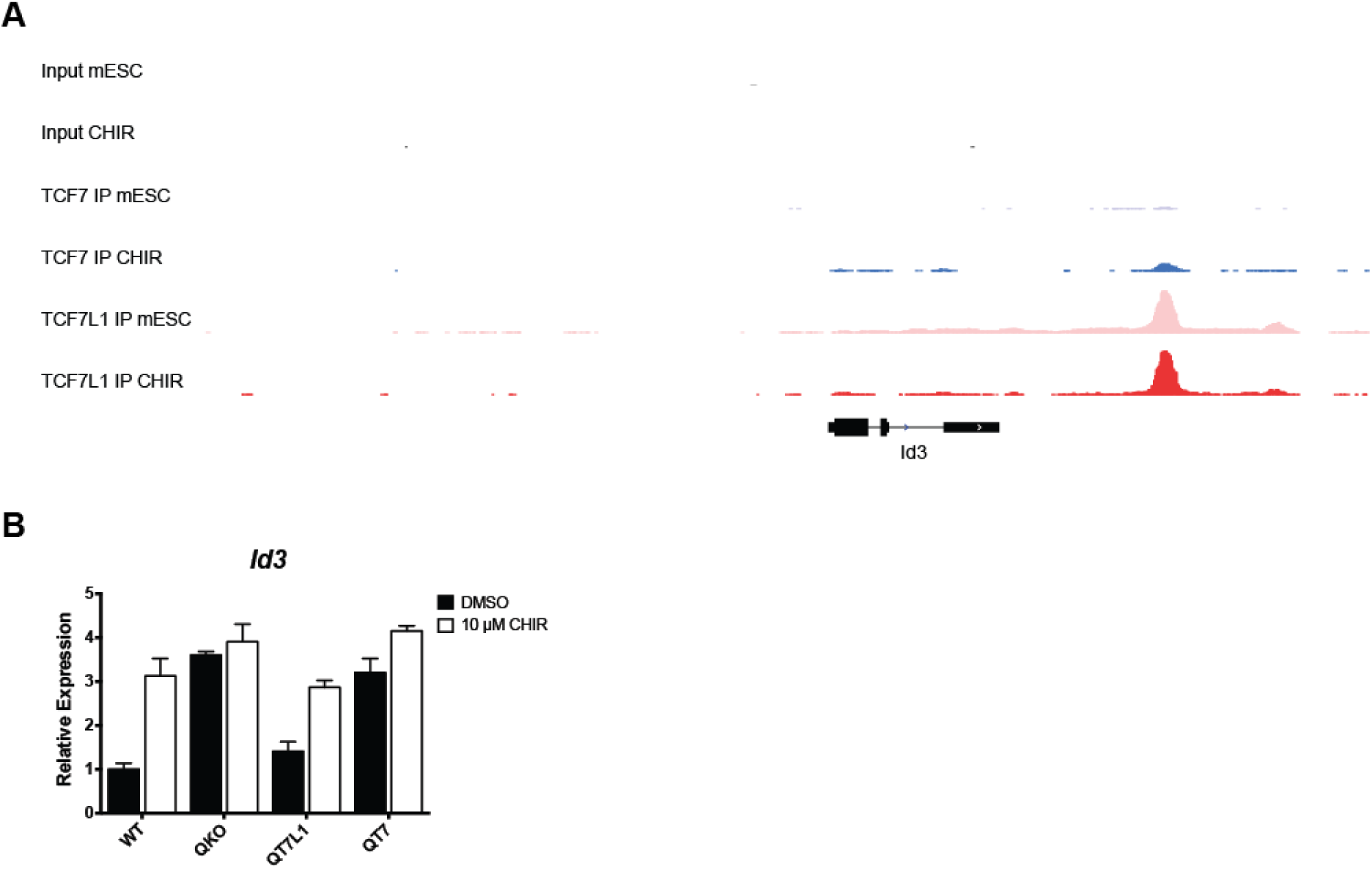
*Id3* activation is mediated by conversion of TCF7L1 into a transcriptional activator in the presence of CHIR. (A) Genomic tracks showing 3xFLAG-TCF7 (blue) and 3xFLAG-TCF7L1 (red) peaks at *Id3* in mESCs cultured in standard medium containing LIF and serum (light shade) or treated with 5 µM CHIR for 14 h (dark shade). Pooled control and CHIR input peaks are in grey and dark grey, respectively. Genomic positions reflect NCBI mouse genome build mm10. (B) qRT-PCR analyses of relative *Id3* transcript levels in WT, QKO, QT7 and QT7L1 cells treated with 10 µM CHIR or DMSO control for 48 h. Bars = mean of three independent experiments ±SEM, normalized to vehicle-treated WT cells.

To further explore this possibility, we examined the effects of CHIR on *Id3* transcript levels in wildtype mESCs, mESCs lacking all 4 full-length TCF/LEFs (QKOs), as well as QKOs rescued with a single copy of 3xFLAG-TCF7L1 (QT7L1) or 3xFLAG-TCF7 (QT7) (Figure 6B). We supplemented standard medium with 10 µM CHIR or DMSO vehicle control for 48 hours. In wildtype mESCs, addition of CHIR caused a 3-fold upregulation of *Id3* transcript levels compared to DMSO control, revealing that *Id3* is responsive to activation by GSK-3 inhibition (Figure 6B). QKO mESCs demonstrated elevated levels of *Id3* in DMSO controls, linking control of its expression to the TCF/LEFs (Figure 6B). Re-expression of a single copy of 3xFLAG-TCF7 did not significantly affect *Id3* expression in QKO mESCs, likely a result of a lack of its binding to the *Id3* locus (Figure 6B). By contrast, rescue with a single copy of 3xFLAG-TCF7L1, restored basal levels of *Id3*, suggesting that in the absence of CHIR, TCF7L1 mediates the repression of *Id3* (Figure 6B). In response to CHIR, QT7L1 cells, in which only 3xFLAG-TCF7L1 is present, demonstrated an upregulation of *Id3* (Figure 6B). Coupled with our observation that TCF7L1 remains bound to the *Id3* locus in cells treated with CHIR, this suggests that CHIR converts TCF7L1 from a repressor of *Id3*, into an activator.

## Discussion

Our findings suggest that in pluripotent mESCs, in both the on and off states of the Wnt/β-catenin signaling pathway, there is minimal overlap between TCF7L1 and TCF7 chromatin occupancy. Employing cell lines with endogenously epitope-tagged TCF7 and TCF7L1 and a well-characterized FLAG antibody in our study offers multiple advantages. The FLAG M2 monoclonal antibody has been extensively used in over 3000 peer-reviews publications (sigmaaldrich.com) and has been used previously for ChIP-seq studies, including a recent study proposing its applicability for large scale ChIP-seq analyses of endogenously tagged transcription factors(27). By using single well-characterized monoclonal antibody to detect endogenously tagged TCF7 and TCF7L1, we are able to conduct quantitative comparisons between these factors. Few studies have examined the genome-wide binding profiles of two TCF/LEF factors simultaneously. Strikingly, we observed minimal overlap between genes bound by TCF7 or TCF7L1 in both standard maintenance conditions and in the presence of the GSK-3 inhibitor, CHIR. This differential binding occurs even though the *de novo* consensus DNA binding motif generated for both factors is a conventional TCF/LEF-binding WRE. In a similar study to ours, Wallmen *et al.* described differential binding of ectopically expressed human TCF7 and TCF7L2 to select Wnt target genes in mESCs including: *Axin2*, *Cdx1*, *T* and *Sp5* (10). Another recent ChIP-seq study in mESCs, using commercially available antibodies to TCF7L1 and TCF7, also revealed minimal overlap between both factors but identified disparate TCF7 and TCF7L1 DNA binding motifs (21).

Wnt-mediated reinforcement of self-renewal and pluripotency has been suggested to involve a switch between repressive TCF7L1 and activating TCF7 through a β-catenin-dependent mechanism (15). Our data suggest that in response to CHIR, not only do β-catenin/3xFLAG-TCF7L1 interactions increase, but 3xFLAG-TCF7L1 is also found bound to a greater number of genes than 3xFLAG-TCF7. While we observed some overlap between TCF7L1 and TCF7 occupancies, there were few regions in which CHIR led to a reduction in TCF7L1 binding with a concomitant increase in TCF7, with the exception of *Axin2* and *T*. Taken together, our data reveal that a TCF7L1-TCF7 switch is not a global requirement for TCF/LEF target gene regulation, but may facilitate Wnt regulation of *Axin2* and *T*.

Our 3xFLAG-TCF7 ChIP-seq data revealed a relatively small number of genes bound in control and CHIR conditions. This is likely a reflection of significantly lower 3xFLAG-TCF7 protein levels, compared to TCF7L1 levels, in these conditions. However, having identified the TCF/LEF consensus binding sequence as our most enriched *de novo* motif for 3xFLAG-TCF7, we are confident that our 3xFLAG-TCF7 data are biologically relevant.

Despite being expressed at low levels, 3xFLAG-TCF7 was more abundant than 3xFLAG-TCF7L1 at some specific Wnt-associated genes, namely: *Zfp703*, *Tcf7*, *Lef1* and *Nkd1*. Presumably, 3xFLAG-TCF7 outcompetes 3xFLAG-TCF7L1 for binding, even though 3xFLAG-TCF7L1 protein is more abundant. Indeed, so called ‘E-tail’ isoforms of TCF7 and TCF7L2, contain a cysteine-clamp that binds to a secondary DNA motif referred to as a ‘helper site’, which is essential for TCF7/TCF7L2-mediated repression and Wnt responsiveness (23, 28, 29). We observed multiple helper sites associated with these particular Wnt target genes. However, there were 3xFLAG-TCF7 and 3xFLAG-TCF7L1 co-occupied Wnt target genes, such as *Axin2*, *Lef1* and *1700001O22Rik* in control conditions and *Apod*, *Zfp36l1*, *mir8110*, *Zfp423*, *mir7020*, and *Sp5*, in CHIR treated cells only.

Some genes containing both WREs and helper sites were found bound to only 3xFLAG-TCF7 or 3xFLAG-TCF7L1 exclusively. This suggests that there are contexts in which TCF7 is unable to outcompete TCF7L1 for binding. In support of this notion, in mouse hair follicle stem cells, TCF7L1 and the E-tail containing TCF7L2, demonstrated a significant amount of overlap in co-bound genes (30). Mechanistically, this could be attributed to suboptimal helper site configurations at specific Wnt target genes (28).

The genomic binding profile of TCF7L1 was the first of the TCF/LEFs to be characterized in mESCs, demonstrating a large overlap in binding with master transcriptional regulators of pluripotency, namely POU5F1, SOX2, and NANOG (11, 12). In our study, in both control and CHIR culture conditions, the second most enriched motif identified was a POU5F1/SOX2/NANOG motif, associated with peaks cobound by all 3 factors. Interestingly, this motif represented a significant proportion of 3xFLAG-TCF7L1 peaks in our control pluripotency maintenance conditions, which increased in the presence of CHIR. Still, we observed minimal overlap between our 3xFLAG-TCF7L1-bound genes in control conditions, and those identified in similar condtions by Marson *et al.* (12). This discrepancy between our study and that of Marson *et al.* could be due to differences in the parental mESCs used in the two studies, differences between the antibodies used for ChIP and/or differences in the chromatin fragmentation methods that were used (sonication vs. enzymatic).

It has been suggested that Wnt signaling regulates TCF7L1 expression through β-catenin mediated removal of TCF7L1 from the chromatin, and subsequent proteasomal degradation (17). Unexpectedly, despite lower 3xFLAG-TCF7L1 levels, we observed only a slight decrease in total 3xFLAG-TCF7L1-bound genes in the presence of CHIR compared control conditions. We also identified nearly an equal number of genes preferentially bound by TCF7L1, with a 43% overlap, between conditions. Our data indicate that 3xFLAG-TCF7L1 is far more abundant than 3xFLAG-TCF7 in CHIR-treated mESCs, and that more 3xFLAG-TCF7L1 than 3xFLAG-TCF7 is associated with β-catenin in this condition. This suggests that TCF7L1 is the primary mediator of Wnt target gene regulation in mESCs and TCF7L1 dissociation from the chromatin is not necessarily required. However, at specific loci such as *Axin2* this mechanism of TCF7L1 derepression is likely employed.

In the absence of a Wnt signal, TCF7L1 is thought to mediate the repression of pluripotency associated genes, including *Nanog* (9, 11–13). Neither 3xFLAG-TCF7L1 nor 3xFLAG-TCF7, negatively correlated with expression of NANOG in self-renewing or differentiating mESC populations, as assessed by flow cytometry and immunofluorescence. In addition, the only ChIP-seq peaks for 3xFLAG-TCF7L1 that we detected at the *Nanog* locus indicated low-level binding to an upstream *Nanog* enhancer that has been shown to be dispensable for pluripotent mESC self-renewal (31). Taken together, our data suggest that TCF/LEFs may mediate indirect effects on *Nanog* expression in a context-dependent manner.

Wnt stimulation of mESCs promotes self-renewal and pluripotency, primarily through the derepression of TCF7L1 (14, 15). This is thought to be promoted by degradation of TCF7L1 upon β-catenin binding (17). Unexpectedly, we observed equal or greater TCF7L1 occupancy, in CHIR-treated mESCs, at many Wnt and pluripotency-related genes. Previously, we identified *Id3* as one of the most upregulated genes in cells lacking all full-length TCF/LEF factors (Moreira et al., 2017). In our current study, we provide data supporting the conversion of TCF7L1 from a repressor into an activator of *Id3* expression in response to GSK-3 inhibition with CHIR. In support of this, in *Xenopus* embryos and mouse keratinocytes, TCF7L1 has been shown to activate a TCF reporter and Klf4, respectively (32, 33).

TCF7 and LEF1 have been classified as activators, whereas TCF7L2 and, especially, TCF7L1, have been suggested to function as repressors. Our TCF7L1 ChIP-seq data suggest that TCF7L1 is the most abundant TCF factor associated with mESC chromatin, mediating the majority of Wnt/β-catenin transcriptional responses. Future experiments are needed to evaluate the roles of the other two TCF/LEF factors, LEF1 and TCF7L2, in mediating β-catenin-dependent transcription in later stages of mESC differentiation, when the level of expression of these factors becomes high enough to be relevant. Also, the roles of the numerous splice variants of the TCF/LEFs need to be incorporated in future studies employing epitope-tagged cell lines, where quantitative comparisons between the various isoforms can be systematically evaluated.

Collectively, the data from our study provide new insights into the mechanisms through which TCF7L1 and TCF7 regulate transcription, highlighted by an unexpected lack of overlap between TCF7L1 and TCF7 binding sites at the whole-genome level and revealing diverse effects of CHIR on the occupancy of both factors.

## Experimental Procedures

Unless otherwise stated, chemicals were obtained from Sigma-Aldrich.

### Cell culture

Routine maintenance of mESCs in serum-containing media, has been previously described (18).

### Chromatin Immunoprecipitation assays

We used the SimpleChIP®Enzymatic Chromatin IP kit according to the manufacturer’s protocol (CST, 9005S). DNA-protein complexes were isolated from 5 x 10E7 mESCs cultured in standard medium, with or without 5 µM CHIR, for 14 h. DNA-protein complexes were crosslinked with 1% paraformaldehyde (11 min, RT). Crosslinking was quenched with 125 mM glycine (5 min, RT). Cells were washed twice with ice-cold PBS and scraped into PBS with 1x protease inhibitor cocktail (PIC). Cells were pelleted and resuspended in buffer A + DTT + PIC, followed by a 10 min incubation on ice. After centrifugation, the nuclear pellet was resuspended in buffer B + DTT + micrococcal nuclease and incubated for 20 min at 37°C with frequent mixing. Chromatin was digested to a length of 150-900 base pairs. 2% inputs were removed and stored overnight at −80°C. Chromatin was immunoprecipitated with anti-FLAG M2 antibody (Sigma-Aldrich, F1804) overnight at 4°C with rotation. DNA-protein-antibody complexes were captured with a 2 h incubation at 4°C using ChIP-grade protein G magnetic beads, followed by 3x low salt and 1x high salt, washes. DNA-protein complexes were resuspended in elution buffer and incubated for 30 min at 65°C. Eluted chromatin was reverse-crosslinked by addition of 0.2M NaCl and proteinase K, followed by incubation at 65°C overnight. DNA was subsequently column purified using supplied reagents and used for quantitative RT-PCR or next generation sequencing.

## Supporting information

## AUTHOR CONTRIBUTIONS

BWD and SM designed the experiments, conducted experiments and wrote the paper. CS, EP and SM conducted experiments. EM provided ChIP-seq analyses and visualization. AB provided guidance with ChIP-seq methodology and analyses.

## ACKNOWLEDGEMENTS

Funding for this study and required infrastructure was provided by the Canadian Institutes of Health Research to BWD (MOP133610), the Canada Research Chairs Program (BWD), the Ontario Ministry of Research and Innovation (BWD), the Canada Foundation for Innovation (BWD), and the OCRiT Project: Ontario Ministry of Economic Development and Innovation (BWD). Bioinformatics analysis was provided by McGill University and Génome Québec Innovation Centre (Montréal, Canada).

