## Supplementary material for "TCF7L1 and TCF7 differentially regulate specific mouse ES cell genes in response to GSK-3 inhibition"

**Supplemental Information**

**TCF7L1 and TCF7 differentially regulate**

**Specific mouse ES cell genes in response to**

**GSK-3 inhibition**

Steven Moreira^1^, Caleb Seo^1^, Enio Polena^1^, Sujeivan Mahendram^1^, Eloi Mercier^2^, Alexandre Blais^3^ and Bradley W. Doble^1*^

**SUPPLEMENTAL FIGURE LEGENDS**

**Figure S1, Related to Figure 1**

(A) Western blot analysis of WT, and both 3xFLAG- TCF7L1 and 3xFLAG-TCF7 clones (A and B), cultured in 15% FBS ± LIF for 48h. Lysates were probed with antibodies against TCF7L1, TCF7, FLAG and β-Tubulin (loading control), as indicated. (B) Quantitative ChIP using the FLAG antibody on chromatin isolated from WT, and both 3xFLAG- TCF7L1 and TCF7 clones, cultured in 15%FBS + LIF for 48h. Percent input was calculated for regions bound by TCF/LEFs in *Axin2* and *Cdx1*, as well as a negative control locus 11kb upstream of *Axin2*. Bars represent the mean of 3 independent experiments ± SEM.

**Figure S2, Related to Figure 2**

(A) Representative flow cytometry profiles of intracellular flow analysis of NANOG, and FLAG levels in 3xFLAG-TCF7L1 and 3xFLAG-TCF7 mESCs (clones not presented in paper) cultured in 15% FBS + LIF (Day 0) and 15% FBS (Day 1-3), as indicated. (B) Graph of the proportion of single-positive FLAG-NANOG+ and double-positive FLAG+NANOG+ cells in WT, 3xFLAG- TCF7L1 and 3xFLAG-TCF7 mESCs. Bars represent the mean of 3 independent experiments ± SEM. (C) Graph of the median fluorescence intensity of NANOG in single-positive FLAG-NANOG+ and double-positive FLAG+NANOG+ cells in WT, 3xFLAG- TCF7L1 and 3xFLAG-TCF7 clones. Bars represent the mean of 3 independent experiments ± SEM. (D) Immunofluorescence analysis of 3xFLAG-TCF7L1 and 3xFLAG-TCF7 mESCs (clones not presented in paper), cultured in 15%FBS + LIF (Day 0) and 15%FBS (Day 1-3), as indicated. Cells were stained for NANOG, FLAG and DAPI. Scale bar represents 50μm. White arrows indicate cells with elevated levels of FLAG and NANOG.

**Figure S3, Related to Figure 3**

(A) Western blot analysis of WT, 3xFLAG- TCF7L1 (clones A and B) and TCF7 (clones A and B) mESCs, cultured in 15% FBS + LIF treated with 5μM CHIR or DMSO control for 14 hours. Lysates were probed with antibodies against TCF7L1, TCF7, FLAG and β-Tubulin (loading control), as indicated. (B) Immunofluorescence analysis of WT, 3xFLAG- TCF7L1 and 3xFLAG-TCF7 mESCs (clones not presented in paper), cultured in 15%FBS + LIF treated with 5μM CHIR or DMSO control for 14 hours, as indicated. Cells were stained for β-catenin, FLAG and DAPI. Scale bar represents 20μm. (C) Quantitative ChIP using the FLAG antibody on WT, and all clones of 3xFLAG- TCF7L1 and 3xFLAG-TCF7 mESCs, cultured in 15%FBS + LIF treated with 5μM CHIR or DMSO control for 14 hours. Percent input was calculated for regions bound by TCF/LEFs in *Axin2* and *Cdx1*, as well as a negative control locus 11kb upstream of *Axin2*. Bars represent the mean of 3 independent experiments ± SEM.

**Figure S4, Related to Figure 4**

(A) Venn diagram showing the overlap between 3xFLAG-TCF7L1-bound genes in mESCs maintained in standard LIF + Serum (mESC) or 14h 5μM CHIR (CHIR) media and TCF7L-bound genes from Table S2 of the supplemental data found in Marson *et al.* (B) Venn diagram showing the overlap between 3xFLAG-TCF7L1 (LEFT) or 3xFLAG-TCF7 (RIGHT) bound genes in mESCs maintained in standard LIF + Serum (mESC) or 14h 5μM CHIR (CHIR) media. (C) Venn diagram showing the overlap between 3xFLAG-TCF7L1- and 3xFLAG-TCF7-bound genes in mESCs maintained in LIF + Serum (mESC) or 14h 5μM CHIR (CHIR) media.

**Figure S5, Related to Figure 5**

(A) Genomic tracks showing 3xFLAG-TCF7 (blue) and 3xFLAG-TCF7L1 (red) peaks at additional Wnt associated genes in mESCs cultured in standard LIF + serum (light shade) or treated with 5μM CHIR for 14 hours (dark shade). Pooled LIF + serum and 14hr 5μM CHIR input peaks are in grey and dark grey, respectively. Genomic positions reflect NCBI mouse genome build mm10.

Figure S1, related to Figure 1

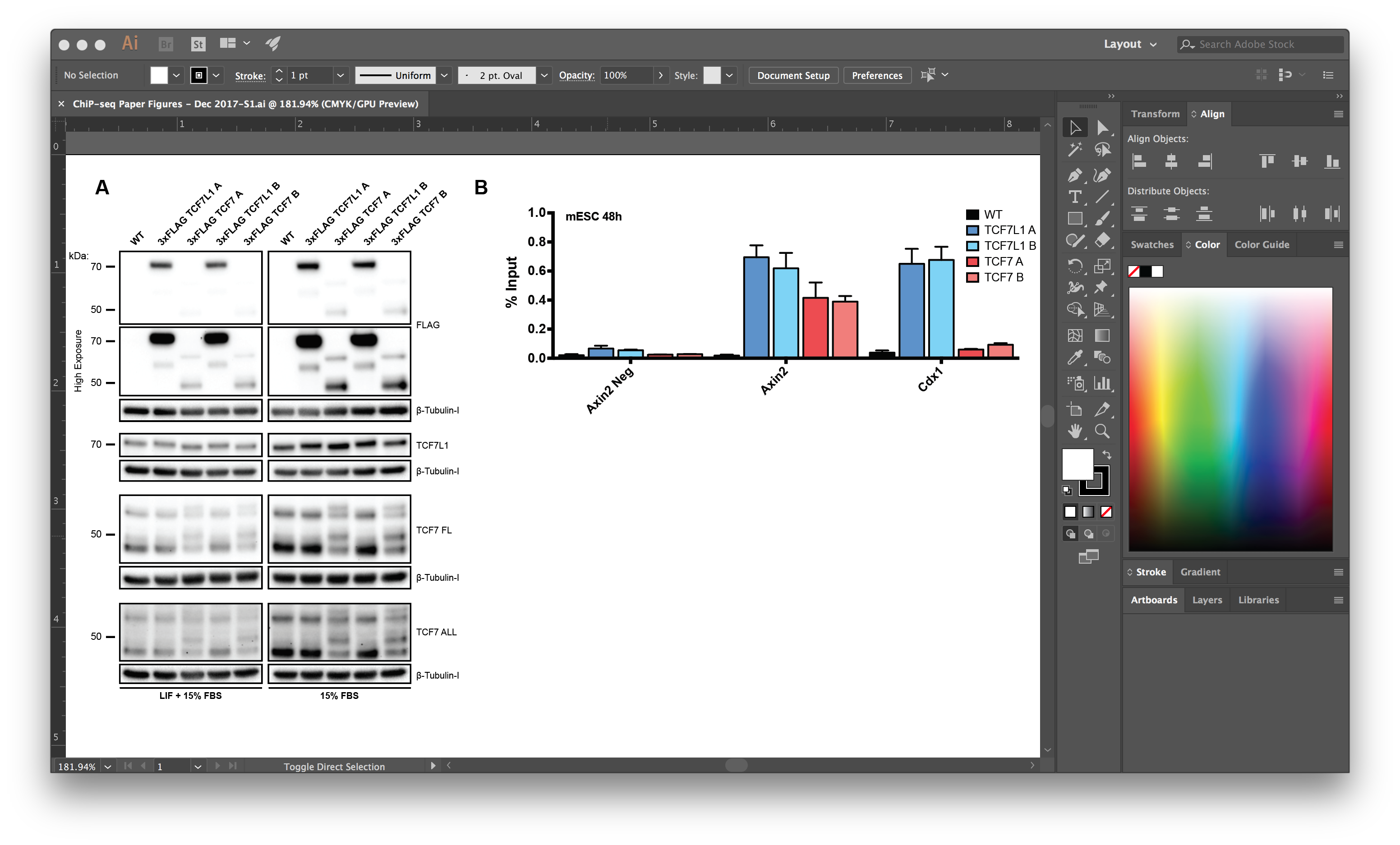

Figure S2, related to Figure 2

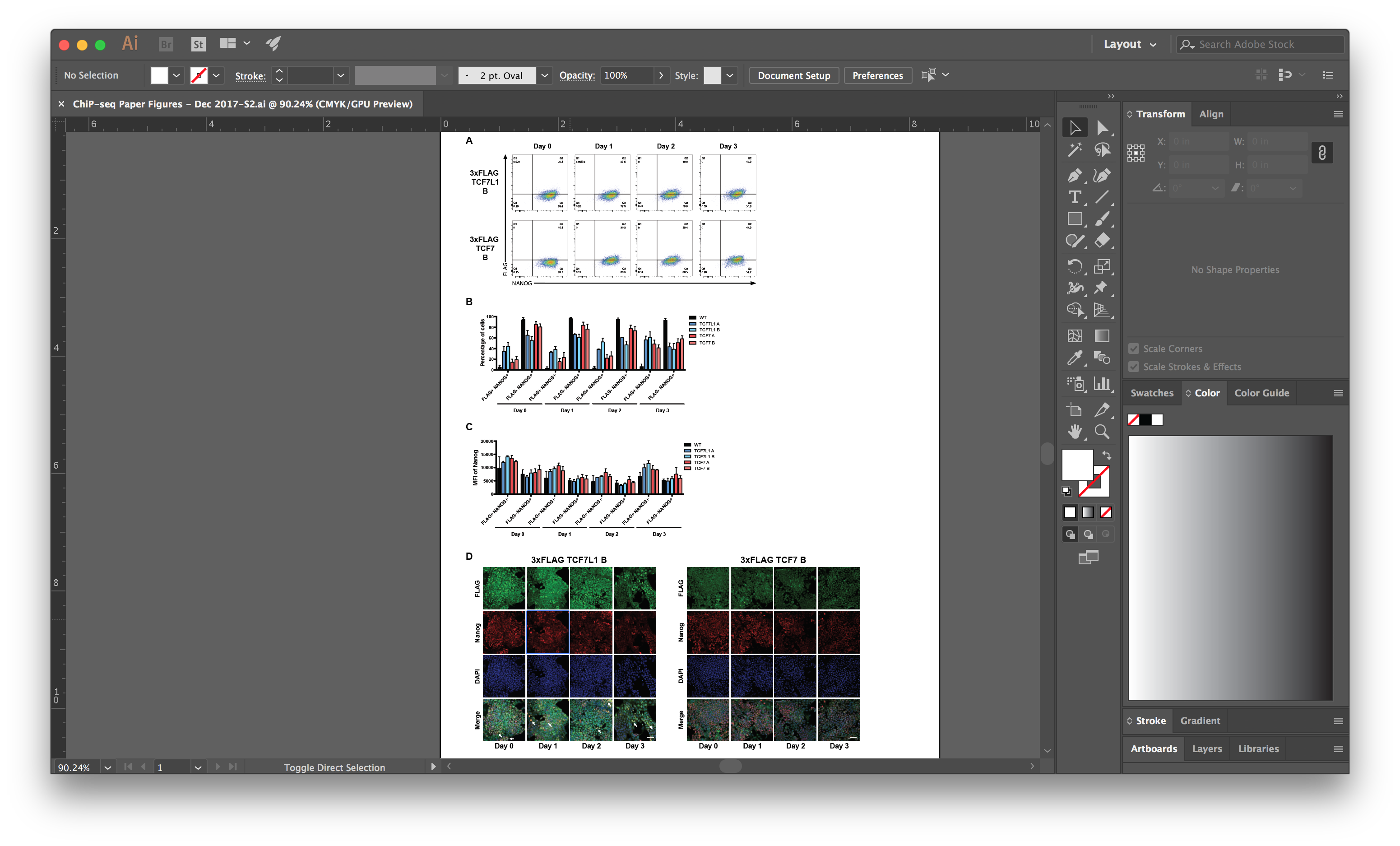

Figure S3, related to Figure 3

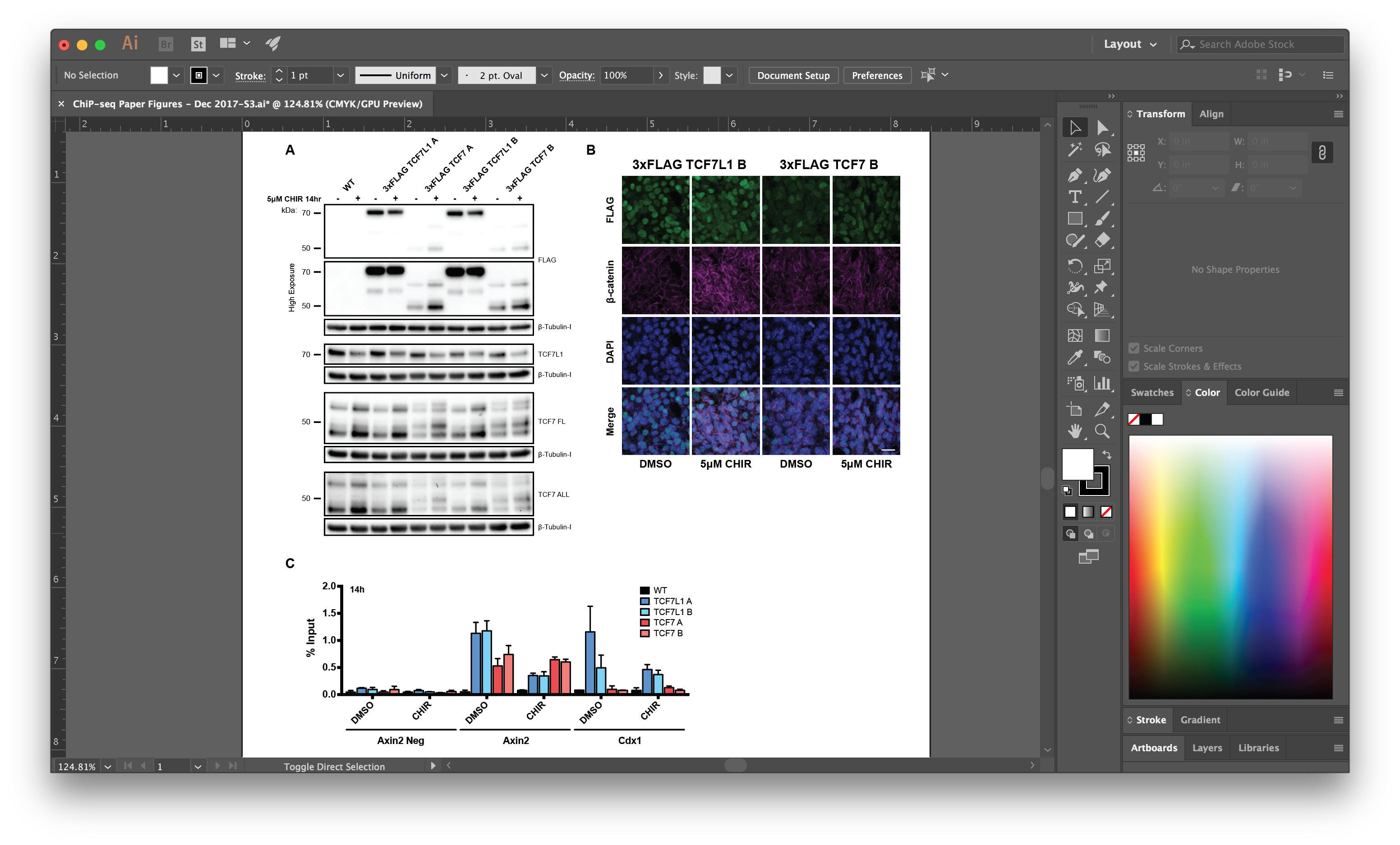

Figure S4, related to Figure 4

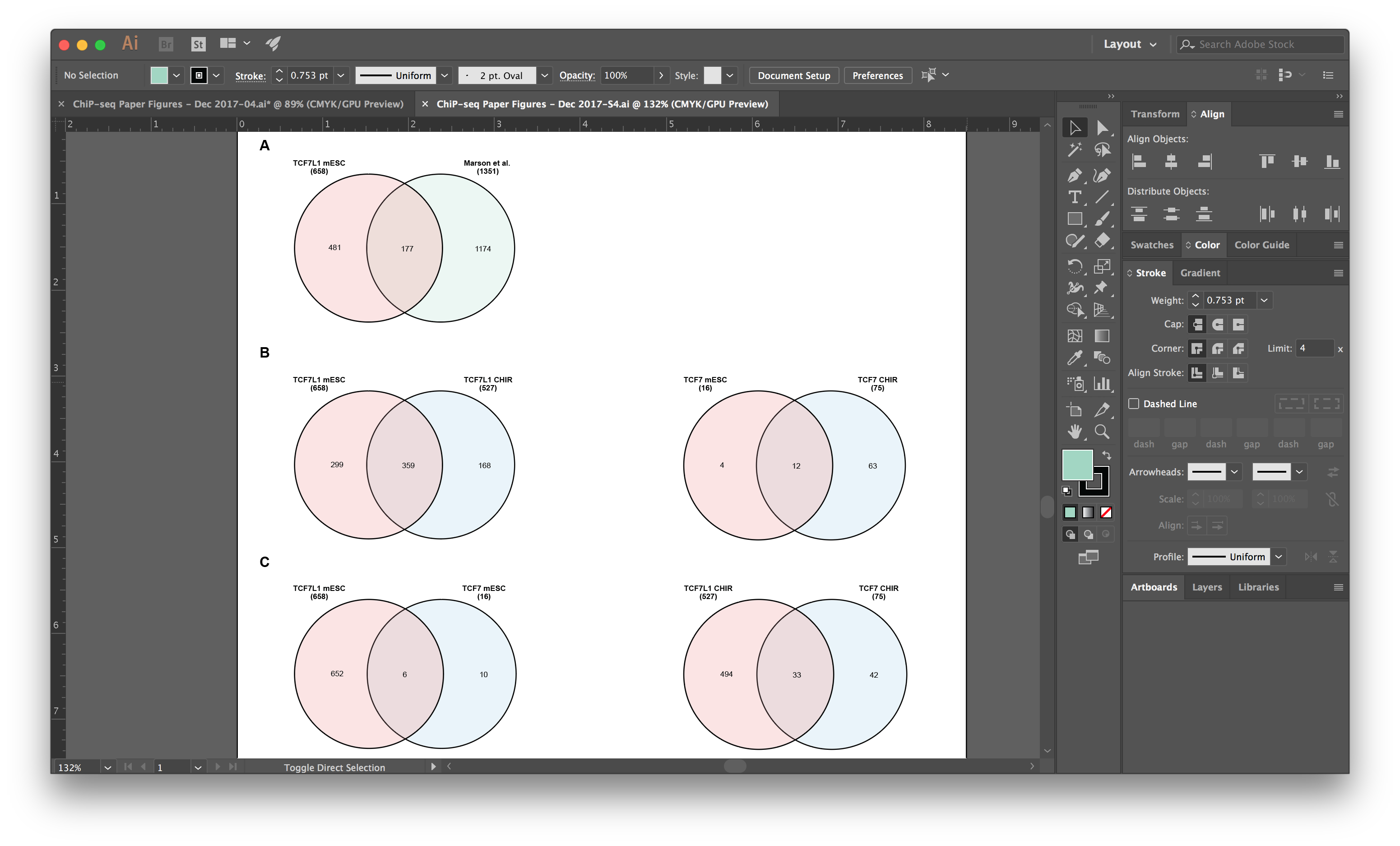

Figure S5, related to Figure 5

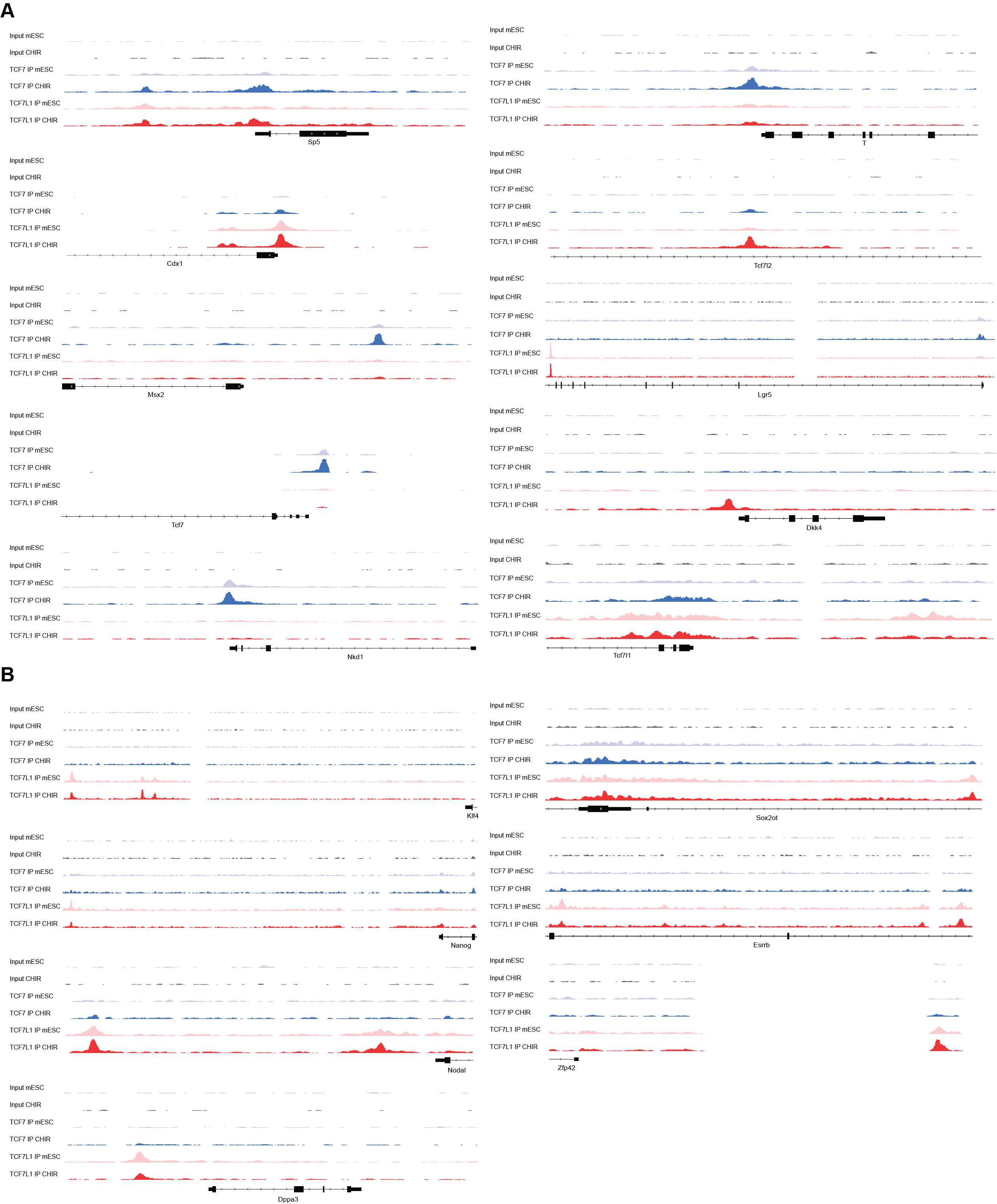

**Table S1 (related to Figure 1)**

Table showing the relative levels of TCF7, TCF7L1 and FLAG in WT compared to 3xFLAG-TCF7 and 3xFLAG-TCF7L1 mESCs cultured in medium containing serum and LIF or serum lacking LIF. The lanes and bands within the western blot were detected using BioRad’s ImageLab™. The relative intensities were calculated by taking the sum of the intensities for all bands in a given lane. The relative intensities for both clones (A and B) were averaged for both TCF7 and TCF7L1. These values were used to calculate the levels of TCF7 and TCF7L1 in WT compared to 3xFLAG-TCF7 and 3xFLAG-TCF7L1, respectively. The average intensity of both clones was also used to calculate the relative levels of TCF7 compared to TCF7L1 in the FLAG blots for both conditions.

**Table S2 (related to Figure 3)**

Table showing the relative levels of TCF7, TCF7L1 and FLAG in WT compared to 3xFLAG-TCF7 and 3xFLAG-TCF7L1 mESCs cultured in medium supplemented with CHIR or DMSO vehicle control for 14 hours. The lanes and bands within the western blot were detected using BioRad’s ImageLab™. The relative intensities were calculated by taking the sum of the intensities for all bands in a given lane. The relative intensities for both clones (A and B) were averaged for both TCF7 and TCF7L1. These values were used to calculate the levels of TCF7 and TCF7L1 in WT compared to 3xFLAG-TCF7 and 3xFLA--TCF7L1, respectively. The average intensity of both clones was also used to calculate the relative levels of TCF7 compared to TCF7L1 in the FLAG blots for both conditions.

**Table S3 (related to Figure 4)**

Table with annotated peaks generated by HOMER from ChIP-seq analyses of WT, 3xFLAG TCF7L1, and 3xFLAG-TCF7 mESCs maintained in standard (LIF+serum) and CHIR (14-hour) media.

**Table S4 (related to Figure 4)**

Table with peaks containing WRE motifs and adjacent helper sites identified by HOMER

ChIP-seq analyses of WT, 3xFLAG TCF7L1, and 3xFLAG-TCF7 mESCs maintained in standard (LIF+serum) and CHIR (14-hour) media.

**Table S5 (related to Figure 5)**

Table with csaw differential binding analysis comparing 3xFLAG-TCF7L1 or 3xFLAG-TCF7 peaks in LIF + serum vs CHIR conditions as well as 3xFLAG-TCF7L1 vs 3xFLAG-TCF7 peaks discovered in LIF + serum or CHIR conditions. logFC.up or logFC.down indicates a gene is higher in the first or second dataset being compared, respectively.

**Supplemental Experimental Procedures**

**Immunofluorescence**

For immunofluorescence staining, cells were washed twice with PBS between all steps. Cells on ibidi^®^ 8-well slides were fixed with cold 4% paraformaldehyde/PBS for 10 minutes at room temperature. Cells were subsequently permeabilized with 0.1% Triton X-100 for 5 minutes at room temperature. Samples were blocked with 10% goat serum/PBS for 1hr followed by incubation overnight at 4°C in primary antibody diluted in 3% goat serum/PBS. The next day, samples were incubated in secondary (Alexa Fluor) antibodies diluted 1:1500 in antibody dilution buffer for 1 hour at room temperature. Cells were mounted with 1-2 drops/well of Pro-Long Gold antifade (containing DAPI) solution (Life Technologies, P36935). Cells were subsequently imaged using a Zeiss LSM 700 laser-scanning confocal fluorescence microscope.

**Intracellular Flow Cytometry**

Cells were plated on 12-well plates. Cells were detached and dispersed into a single-cell suspension with Accutase (Innovative Cell Technologies). After washing once with PBS, cells were resuspended in in PEF buffer (2% FBS and 2 mM EDTA in PBS). Cells were subsequently strained through a 35 μm flow tube and fixed with BD Cytofix™ fixation buffer (554655, BD Biosciences). Cells were washed with 1x BD Perm/Wash™ buffer (554723, BD Biosciences) and pelleted. Cells were then incubated in anti-Nanog and FLAG antibodies diluted (1/800) in 1x BD Perm/Wash™ buffer overnight at 4°C. Cells were then washed with 1x BD Perm/Wash™ buffer and pelleted. Pellets were resuspended in secondary antibodies, goat anti-mouse Alexa Fluor 488 and goat anti-rabbit Alexa Fluor 647, diluted (1/4000) in 1x BD Perm/Wash™ buffer, and incubated for 1h at 4°C. A final wash was performed with 1x BD Perm/Wash™ buffer and cells were pelleted. Cells were subsequently resuspended in PEF buffer and analyzed by flow cytometry.

**Cell lysate preparation**

Cell lysates were prepared as previously described (Moreira et al., 2017).

**Western Blot Analysis**

Western blots were performed as previously described (Moreira, et al., 2017).

**Co-Immunoprecipitation Assays**

Approximately 5-10 x 10^6^ cells of the indicated mESC lines were cultured on 100 mm gelatin-coated dishes in standard mESC medium, prior to initiation of the assay. The cells were rinsed twice with PBS and lysed for 10 minutes on ice with Buffer LB (14321B, ThermoFisher Scientific), which contained 100 mM NaCl, 0.5% Triton™ X-100, 2 mM MgCl_2_, 1 mM DTT, 1x Halt Protease and Phosphatase Inhibitor cocktail (78440, ThermoFisher Scientific). The cells were subsequently harvested, and the lysates were clarified at 18 000 x g for 10 minutes. The supernatant was transferred to a new pre-chilled tube and the lysates were quantified by the Lowry method (DC Protein Assay; 5000112, Bio-Rad). Equivalent amounts of protein extract were used per IP within each co-immunoprecipitation experiment (typically ~500μg of protein per IP). The following antibody was used for the assay, mouse anti-β-catenin (610153, BD Transduction). 7.5 μg per IP was pre-conjugated to 1.5mg of Protein-G Dynabeads™ M-270 Epoxy for 24 hours at 37°C according the manufacturer’s instructions (14321B, ThermoFisher Scientific). Extracts from the indicated mESC lines were added to the antibody-bead complex, and the mixture was incubated for 1 hours at 4°C, rotating. The immunoprecipitates were subsequently rinsed 3 times using the lysis buffer (including inhibitors) followed by a final wash with LWB buffer (14321B, ThermoFisher Scientific). Samples were eluted from the beads with EB buffer (14321B, ThermoFisher Scientific) via a 5-minute incubation at 25°C. Prior to western blot analysis, 3x NuPage™ LDS buffer (NP0007, ThermoFisher Scientific) with 5% TCEP Bond Breaker solution (77720, ThermoFisher Scientific) was added to the eluate without boiling.

**Quantitative RT-PCR**

Total RNA was isolated using the PureLink RNA Mini Kit (Life Technologies) and 1 μg was used to generate cDNA with qScript cDNA SuperMix (Quanta Biosciences). The cDNA was diluted 1/5 to a final volume of 100 μL and 3 μL of this was used for each 10 µL PCR reaction with SsoAdvanced SYBR Green SuperMix (BioRad). *Rpl13a* was used as the reference gene. Bio-Rad’s CFX96 instrument and software were used to determine relative gene expression levels using the delta-delta Ct method. Primer sequences were designed using IDT’s online primer design software (idtdna.com) or were obtained from prior publications. For experiments in which transcript levels were assessed following CHIR99021 treatment of WT ESCs, ESCs were plated into LIF- containing medium supplemented with CHIR99021 at 10 μM for 48 hours before RNA isolation.

**Antibodies**

The following primary antibodies were used for western blot, co-immunoprecipitation, and/or immunofluorescent staining: mouse anti-β-Tubulin-I (T7816, Sigma); mouse anti-FLAG M2 (F1804, Sigma); rabbit anti-TCF7L1 (ab86175, Abcam); rabbit anti-TCF7 FL (C63D9, Cell Signaling Tech.); rabbit anti-TCF7 All (C46C7, Cell Signaling Tech.); rabbit anti-Nanog (A300-397A, Bethyl Laboratories); mouse anti-β-catenin (610153, BD Transduction); rabbit anti-Non-phospho-β-catenin (D13A1, Cell Signaling Tech.); and rabbit anti-β-catenin Amino-terminal antigen (9581S, Cell Signaling Tech.).

Fluorochrome-conjugated secondary antibodies were obtained from ThermoFisher Scientific and used at 1/1500 (Molecular Probes): goat anti-mouse Alexa Fluor 488; and goat anti-rabbit Alexa Fluor 647.

**Generation of 3xFLAG- *TCF7L1* and TCF7 knock-in mESC lines using TALENs.**

*TCF7L1* and *TCF7* targeting constructs and TALENs were designed and assembled as previously described (Moreira, et al., 2017)*.* Cell lines were generated as previously described (Moreira, et al., 2017). Two independent clones of both 3xFLAG-TCF7L1 and 3xFLAG-TCF7 knock-in mESCs were used for subsequent analyses. Heterozygosity was confirmed using western blotting, Sanger sequencing and genomic DNA genotyping.

**ChIP-seq library preparation and next generation sequencing**

Library preparation and sequencing was performed by McMaster University’s Farncombe Metagenomics Facility. ChIP DNA was initially quantitated using the QuantiFluor dsDNA System reagents (Promega, E2670) and the Promega QuantiFluor ST fluorometer. ChIP-seq libraries were prepared using NEBNext^®^ Ultra™ II DNA library preparation kit for Illumina^®^ according to manufacturer’s instructions (New England Biolabs, E7645). Input libraries from WT, 3xFLAG- TCF7L1 clones A and B, and 3xFLAG-TCF7 clones A and B, cultured in standard mESC medium containing LIF and serum (L+S) or the same medium supplemented with 5μM CHIR for 14h, were pooled to create a L+S and CHIR inputs, respectively. Fragment size distribution for all libraries was assessed using a Bioanalyzer and concentrations were assessed using qPCR on BioRad’s iCycler IQ5 using the KAPA-SYBR Fast qPCR Master Mix (BioRad, KM4105). Sequencing was configured for single-end 51 bp reads and 8 bp dual indices on an Illumina HiSeq 1500 instrument, with Rapid v2 chemistry and on-board cluster generation. Input libraries from L+S and CHIR, were pooled with libraries generated from anti-FLAG ChIPs conducted on chromatin isolated from WT, 3xFLAG-TCF7L1 clones A and B, and 3xFLAG-TCF7 clones A and B, cultured in L+S or CHIR and were sequenced across 4 lanes, resulting in approximately 50 million reads per sample.

**ChIP-seq trimming, alignment, and tracks**

Reads were trimmed from the 3' end to have a phred score of at least **30**. Illumina Universal adapters were removed from the reads. Resulting reads shorter than 50 bp and orphaned reads were discarded. Trimming and clipping were performed using Trimmomatic v0.35 (Bolger et al., 2014). Trimmed reads were then aligned to the mouse reference genome GRCm38 using BWA v0.7.12 (Li et al., 2009). Multi-mappers and duplicate reads were excluded from subsequent analyses. Tracks were generated using bedtools v2.27.0 (Quinlan and Hall, 2010). Bedgraphs were generated from bam files and normalized by a scale factor, which is inversely proportional to the number of reads (genomeCoverageBed -bg -split -scale $scalefactor).

**ChIP-seq peak calling, annotation, motif discovery, and gene ontology**

Narrow peaks were called by MACS v2.1.0 (Zhang et al., 2008) in single end mod (format = BAM), no shifting model (-nomodel) and defaults parameters (mfold = [5,50], q-value cut-off = 0.05). Input-CHIR-Pool and Input-mESC-Pool were used as control for CHIR and mESC samples respectively. Peak annotation, known and *de novo* motif discovery, and gene ontology analysis were performed by HOMER v4.7 (Heinz et al., 2010). Peak tracks were visualized using the Integrative Genomics Viewer (IGV) v2.4 (Robinson et al., 2011).

**Differential Binding Analysis**

Differential binding analysis was performed with the R packages csaw v1.10 (Lun and Smyth, 2015) and edgeR 3.18.1 (Robinson et al., 2009) with R 3.4.1. Briefly, the windowCounts function was used to count the number of reads in a sliding window in every BAM file with the following parameters: minimum quality of 30, fragment length of 150, single-end reads and a window size of 50. After calculating for normalization factors and filtering with negative control (using wildtype samples) to remove uninterested windows, the remaining regions were tested for differential binding using the edgeR workflow. Benjamini-Hochberg multiple test correction method was used to compute the False Discovery Rate within each pairwise comparison.

**TCF/LEF helper site identification**

To identify potential interactions between TCF/LEF motifs and helper sites, we first identified the presence of the WRE motifs within MACS identified at TCF7 or TCF7L1 peaks, using HOMER. Helper sites were subsequently identified 200 bp up- or downstream of each WRE, using HOMER. Position Weight Matrices (PWM) were manually created for the core binding site of TCF/LEF motifs (5’-CTTTGWWS-3’ (W=A/T S=G/C) and the helper site 5’-RCCGCC-3’ (R=A/G) (Hoverter et al., 2014). HOMER v4.7 (Heinz et al., 2010) was used to scan the previously identified peaks for these motifs with a score of 5 or more. Peaks containing both motifs were selected for further analyses.
